# Structural and biochemical mechanism for increased infectivity and immune evasion of Omicron BA.1 and BA.2 variants and their mouse origins

**DOI:** 10.1101/2022.04.12.488075

**Authors:** Youwei Xu, Canrong Wu, Xiaodan Cao, Chunyin Gu, Heng Liu, Mengting Jiang, Xiaoxi Wang, Qingning Yuan, Kai Wu, Jia Liu, Deyi Wang, Xianqing He, Xueping Wang, Su-Jun Deng, H. Eric Xu, Wanchao Yin

## Abstract

The Omicron BA.2 variant has become a dominant infective strain worldwide. Receptor binding studies reveal that the Omicron BA.2 spike trimer have 11-fold and 2-fold higher potency to human ACE2 than the spike trimer from the wildtype (WT) and Omicron BA.1 strains. The structure of the BA.2 spike trimer complexed with human ACE2 reveals that all three receptor-binding domains (RBDs) in the spike trimer are in open conformation, ready for ACE2 binding, thus providing a basis for the increased infectivity of the BA.2 strain. JMB2002, a therapeutic antibody that was shown to have efficient inhibition of Omicron BA.1, also shows potent neutralization activities against Omicron BA.2. In addition, both BA.1 and BA.2 spike trimers are able to bind to mouse ACE2 with high potency. In contrast, the WT spike trimer binds well to cat ACE2 but not to mouse ACE2. The structures of both BA.1 and BA.2 spike trimer bound to mouse ACE2 reveal the basis for their high affinity interactions. Together, these results suggest a possible evolution pathway for Omicron BA.1 and BA.2 variants from human-cat-mouse-human circle, which could have important implications in establishing an effective strategy in combating viral infection.

## INTRODUCTION

The Omicron variants BA.1 and BA.2 of severe acute respiratory syndrome coronavirus 2 (SARS-CoV-2), the causative virus of COVID-19, are two sister variants of concerns (VOCs) that have infected hundreds of millions of people worldwide.^1^ In particular, the BA.2 strain has overtaken the BA.1 strain to become the current dominant infective strain that continues to rampage across the globe (Fig. 1a).^2^ Both BA.1 and BA.2 strains are evolved independently from predominant VOC strains, including Alpha, Beta, Gamma, and Delta variants.^3^ Mutational profile analysis suggests that the Omicron variants may arise from evolution through mouse as a host.^4^ The biochemical and structural basis for the viral infection to mouse remains largely unknown.

**Fig 1.**
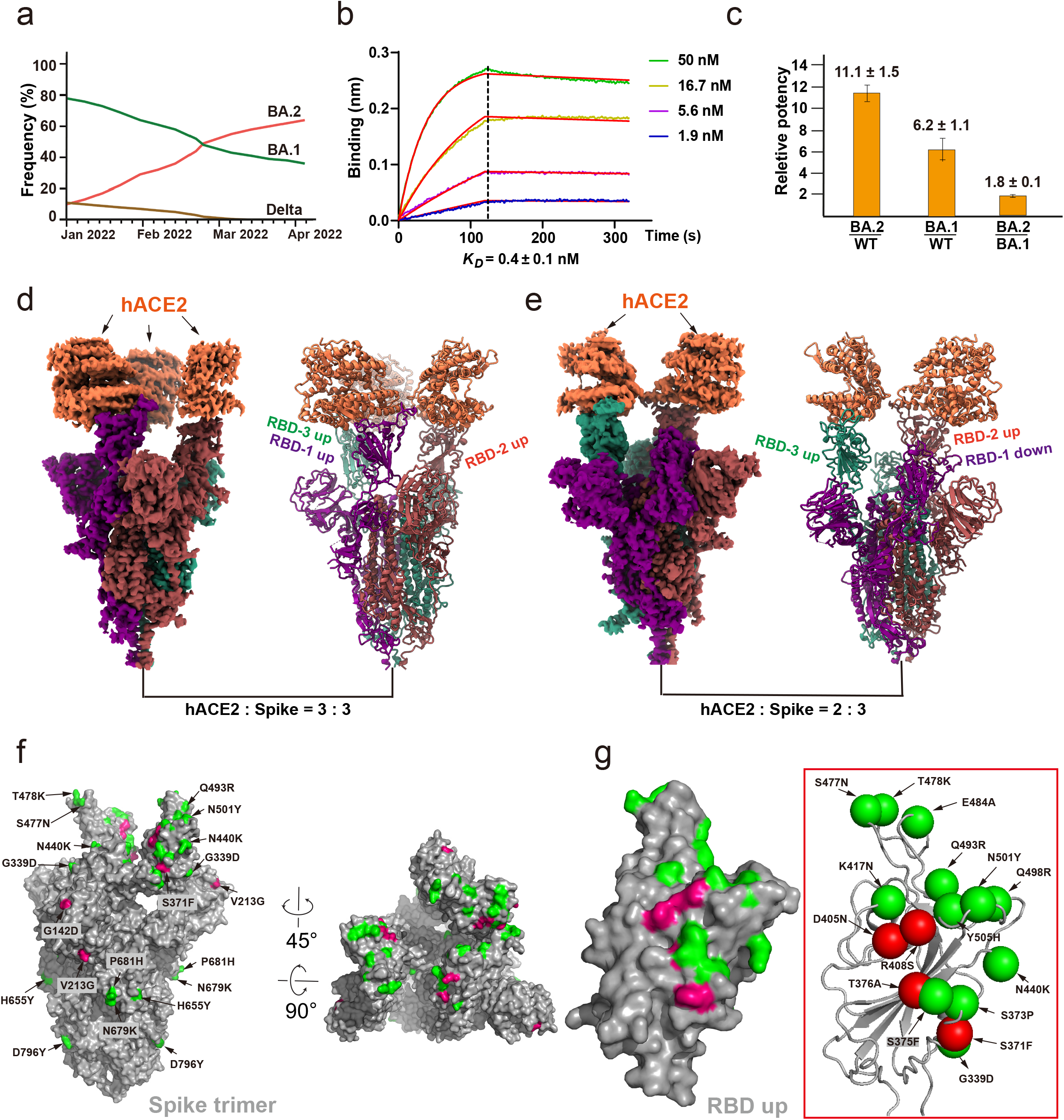
SARS-CoV-2 Omicron BA.2 spike protein with higher affinity to human ACE2. **a**. The infectious frequency of SARS-CoV-2 Delta, Omicron BA.1, and Omicron BA.2 strains since January 2022 to April 2022. **b**. Binding curves of the Omicron BA.2 spike trimer to human ACE2. *K*_D_ values were determined with Octet Data Analysis HT 11.0 software using a 1:1 global fit model. **c**. Relative potency of WT, BA.1, and BA.2 with the ratio of their *K*_D_ values. **d-e**. Cryo-EM density maps of the hACE2-Omicron BA.2 spike trimer complexes with hACE2 and BA.2 spike in 3:3 molar ratio (d) or in 2:3 molar ratio (e). **f**. The locations of Omicron BA.2 mutations on the Spike trimer. Spike trimer is shown in surface. The shared mutations between BA.1 and BA.2 are colored in green and the BA.2 its own mutations are colored in red. **g**. The locations of 16 Omicron BA.2 mutations on the RBD. RBD is shown in surface. The shared mutations between BA.1 and BA.2 are colored in green and the BA.2 its own mutations are colored in red, including S375F, T376A, D405N, R408S, and N440K.

The trimeric spike protein is a major membrane surface glycoprotein of SARS-CoV-2 that mediates the binding to the host receptor ACE2 and subsequent viral entry into cells.^5-7^ The matured spike protein contains two subunits: an ACE2 binding S1 subunit and a membrane-fusion S2 subunit. Within the S1 subunit is an N-terminal domain (NTD) of unknown function and a C-terminal receptor-binding domain (RBD). The S2 subunit contains a conserved fusion peptide (FP) motif that is mostly hydrophobic and capable of mediating viral fusion with host cells.^7^ Extensive structures of the spike trimer reveal the open and closed conformations for the three RBDs within the spike trimer, where the open “up” RBD conformation is prerequisite for ACE2 binding.^3, 6, 8-12^ Upon ACE2 binding, the spike trimer undergoes conformational changes to expose the FP motif of the S2 subunit to initiate membrane fusion that allows the release of the viral genome into host cells.^13^

The spike protein is also the major target of immune response to infection, vaccination, and antibody therapy.^14-18^ Most VOCs such as the Delta variant contain 7-10 mutations in the spike protein. In contrast, Omicron BA.1 and BA.2 have 37 and 31 mutations in their spike protein.^19^ Structural studies of the Omicron BA.1 spike trimer reveal that most of the 37 mutations are mapped on the surface of the spike protein, with many of them enriched in known epitopes of therapeutic antibodies.^3, 20-22^ In addition, the Omicron BA.1 spike protein binds to the human ACE2 with 6-7 fold higher potency than WT spike and the Omicron BA.1 spike trimer is less stable and prone to the open “up” conformation that is ready to interact with ACE2.^3^ These studies have provided the molecular basis for the increased infectivity and immune evasion of Omicron BA.1 to vaccination and antibody therapy. However, the Omicron BA.2 variant is even more contagious and has surpassed BA.1 as the dominant infective strain currently in many regions across the world (Fig. 1a).^2^ The Omicron BA.2 spike protein has 22 amino acid differences from the BA.1 spike protein (Supplementary information, Fig. S1), more than 7 amino acid differences of the Delta variant from the original WT strain.

Although the symptoms of infection by Omicron BA.1 and BA.2 appear to be less severe,^23^ antibody therapy may be essential for critically ill patients. JMB2002, a broad-spectrum therapeutic antibody, is known to inhibit both the WT strain and the Omicron BA.1 variant.^3, 24^ However, the efficacy of JMB2002 against the Omicron BA.2 is unknown. In this paper, we report the cryo-EM structures of the Omicron BA.2 spike trimer in complex with the human ACE2 and JMB2002. We also determined the spike trimer from both BA.1 and BA.2 in complex with the mouse ACE2. Together with comprehensive biochemical studies, our structures provide a basis for the higher transmission and immune evasion of the Omicron BA.2 variant and possible mouse origins of Omicron BA.1 and BA.2 variants.

## RESULTS

### Characterization of the interaction of the Omicron BA.2 spike trimer with hACE2

To study the mechanism for enhanced transmission of Omicron BA.2, we first characterized the interaction between the human ACE2 (hACE2) with the spike extracellular domain trimer from SARS-CoV-2 Omicron BA.2, BA.1, and WT strains, all of which contain mutations in the furin cleavage site and proline substitutions (2P or 6P) to stabilize the prefusion conformation. Dimeric hACE2 bound to immobilized Omicron BA.2 spike trimer with a dissociation constant (*K*_D_) value of 0.4 ± 0.1 nM, which is approximately 11-fold higher than that with WT spike trimer (*K*_D_ =4.7 ± 0.6 nM) and is nearly 2-fold higher than that with BA.1 spike trimer (*K*_D_ =0.75 ± 0.1 nM). (Fig.1b and 1c). We also examined the interactions of monomeric hACE2 with Omicron BA.2 spike trimer with the *K*_D_ value of 3.2 ± 0.7 nM, which is approximately 5-fold higher than that with WT spike trimer (*K*_D_ =15.0 ± 0.6 nM) and is around 2-fold higher affinity than that with BA.1 spike trimer (*K*_D_ =6.4 ± 1.0 nM) (Supplementary information, Fig. S2). The observed interactions between hACE2 and WT spike trimer, and between hACE2 and BA.1 spike trimer were consistent with previously published data.^3^ The enhanced interaction of Omicron BA.2 spike trimer protein with hACE2 may be one of the key factors that contribute to the increased transmissibility of the BA.2 strain.

To gain structural insights into the potent binding of the Omicron BA.2 spike trimer to hACE2, we reconstituted the Omicron BA.2 spike trimer-hACE2 complex with an excess of hACE2, followed by size exclusion chromatography and cryo-electron microscopy (cryo-EM) analysis (Supplementary information, Fig. S3). We observed two distinct structure states of the BA.2 spike trimer-hACE2 complex (Figs. 1d and 1e). In the first state, each spike molecule from the BA.2 spike trimer binds to one hACE2 molecule and all three RBDs are in an open up position for hACE2 binding (3-hACE2-bound BA.2 structure) (Fig. 1d). In the second structure state, two of three RBDs are in the open up position, both of which bind to hACE2 (2-hACE2-bound BA.2 structure). The third RBD is in a clear down position and shows the direct interaction with an up RBD, as it was observed previously (Fig. 1e).^3^ Particle classification reveals that about 42% and 58% spike particles bind with hACE2 in 3:3 and 3:2 molar ratio, respectively, albeit an excess of hACE2 were incubated with spike trimer during sample preparation. Notably, only one hACE2 bound to one RBD from the spike trimer was observed in our previous cryo-EM structural analysis of Omicron BA.1 spike trimer-hACE2 complex.^3^ In contrast, in the Omicron BA.2 spike trimer-hACE2 complex, the spike trimer bound to at least two or three hACE2, indicating a stronger hACE2 binding tendency of the BA.2 spike trimer.

Mapping the 31 mutations onto the Omicron BA.2 spike trimer reveals that 23 mutations are distributed on the surface and 3 are on the interior of the spike trimer (Fig. 1f). 22 out of 31 mutations are different from BA.1, while the rest of the mutations share the same substitutes with BA.1. Similar with 15 mutations in the RBD of BA.1 spike, 16 mutations are located on the RBD domain of BA.2 spike (Fig. 1g), which serves as ACE2 binding and the epitopes for 90% of antibodies.^3, 25-27^ These mutations could cause the immune evasion to vaccines and therapeutic antibodies^28^, however, some antibodies which do not rely on those sites could retain the immunity.

Focus refinement on RBD-hACE2 region resulted a reconstruction map at 3.0 Å resolution (Supplementary information, Fig. S4). The cryo-EM density map was of sufficient quality to enable us to build an atomic model of the RBD-hACE2 structure (Fig. 2a). The overall assembly of the Omicron BA.2 RBD-hACE2 complex closely resembles the Omicron BA.1 RBD-hACE2 complex, where the receptor binding motif (RBM) is identical. Briefly, the side chain of Q493R forms a new salt bridge with E35 of hACE2, Q498R forms a new salt bridge with D38 of hACE2, and K417N loses a salt bridge (Figs. 2b and 2c), these key mutations resulted in an enhanced binding to hACE2. Thermal shift assays showed that the spike trimers from Omicron BA.2, Omicron BA.1, and WT displayed two melting temperatures, which were assigned previously, with the lower Tm for the RBD and the higher Tm for the spike trimer.^3^ The Tm values for the RBD from Omicron BA.2, Omicron BA.1, and WT are 47.4 °C, 44.5 °C, and 52.5 °C (Fig. 2d), respectively, indicating that RBD in Omicron BA.2 is more stable than that from BA.1, but less stable than the WT. In the BA.2 RBD, the substitute of D405, N405 forms two inter-molecular hydrogen-bonds with the side chain of R403 and the main chain of G504 (Fig. 2e). The BA.1 RBD lacks these interactions because the distance between the side chain of D405 and R403 is about 4.5 Å (Fig. 2e). The additional interactions in the BA.2 RBD above could contribute to its higher stability than the BA.1 RBD. The lower Tm2, which corresponds to the dissociation of the BA.2 spike trimer, indicates its more dynamic nature than BA.1 spike trimer (Fig. 2d). The BA.1 and BA.2 spike trimers are different in three additional residues, including T547K in the S1 region, D856K and L981F in S2 region. These mutations are found in the BA.1 spike but not in the BA.2 spike. Among the three different residues, D856K in BA.1 from protomer 1 forms a salt bridge with D571 from the adjacent protomer, which could stabilize the BA.1 spike trimer (Fig. 2e). In BA.2 strain, D856 lacks such interaction, thus, the BA.2 spike trimer shows lower melting temperature under our thermal shift assays.

**Fig 2.**
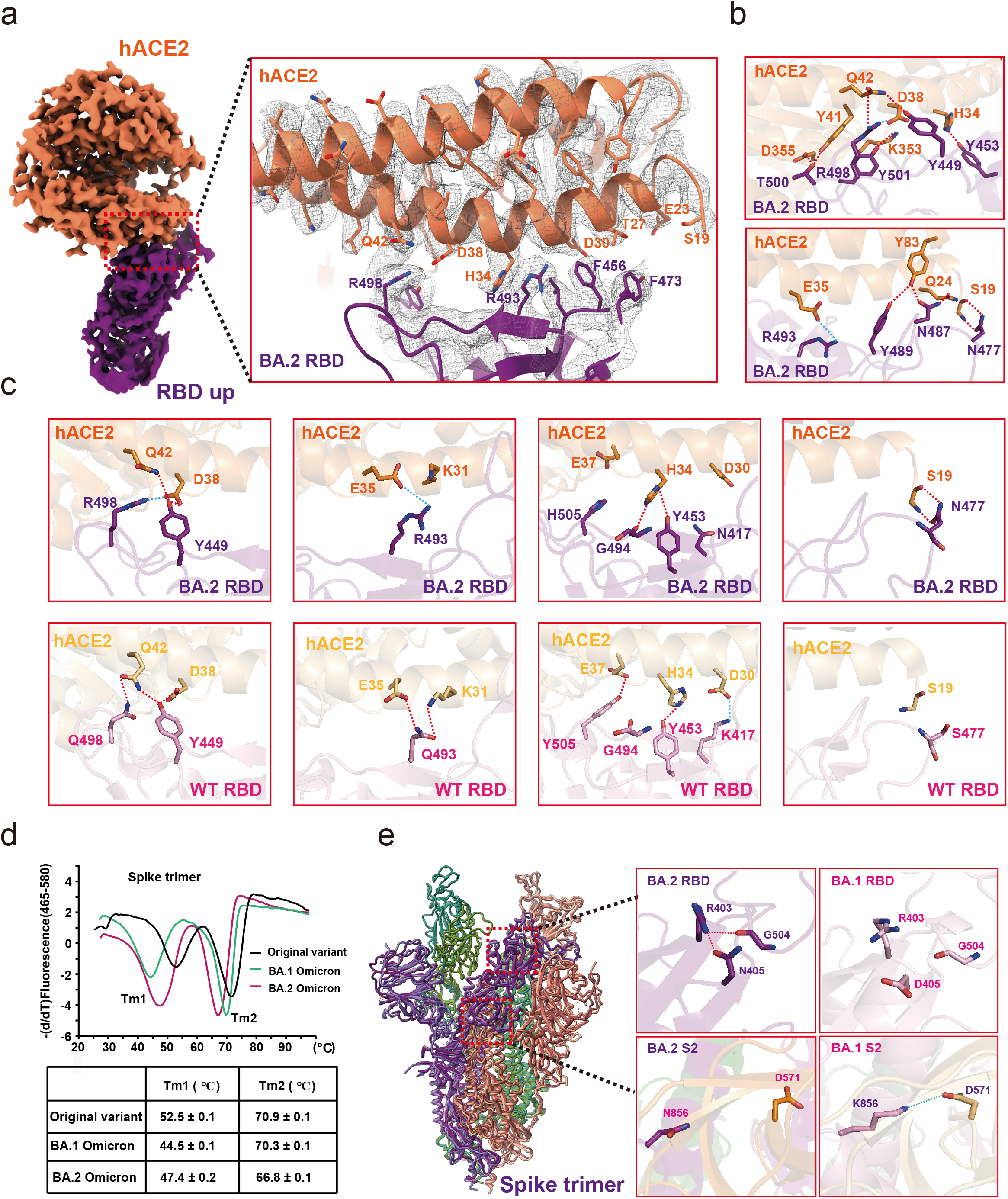
Structural analysis of Omicron BA.2 RBD and hACE2. **a**. Cryo-EM density map of Omicron BA.2 RBD bound to hACE2. Residues are shown in sticks with the correspondent cryo-EM density represented in mesh. hACE2 is colored in coral. The Omicron BA.2 RBD is colored in purple. **b**. Interactions between Omicron BA.2 RBD and hACE2. **c**. Comparison of Omicron BA.2 RBD-hACE2 and WT RBD-hACE2 interfaces. Up panels, Omicron BA.2 RBD-hACE2 with hydrogen bonds and salt bridges interactions. Down panels, WT RBD-hACE2 with hydrogen bonds and salt bridges interactions. Interactions of hydrogen bonds and salt bridges are in dotted lines. **d**. Thermal stability shift analysis of the Omicron BA.2, Omicron BA.1, and WT spike trimer. **e**. The mutation-induced conformation changes of Omicron BA.2 RBD comparing with WT RBD.

### Inhibition of ACE2 binding to the BA.2 spike trimer by an anti-Omicron antibody JMB2002

Previously, we reported the potent neutralization activity of a clinical stage antibody, JMB2002, which could effectively neutralize Omicron BA.1 as well as variants Alpha, Beta, and Gamma, but not Delta.^3, 24^ To evaluate the neutralizing activity of JMB2002 against the Omicron BA.2 variant, we first evaluated the binding of JMB2002 to the Omicron BA.2 spike trimer. JMB2002 Fab recognized the Omicron BA.2 spike trimer with *K*_D_ of approximately 2.6 nM. Meanwhile, JMB2002 IgG bound to the Omicron BA.2 spike trimer with *K*_D_ of approximately 0.3 nM (Fig. 3a). The potency of JMB2002 to the Omicron BA.2 spike trimer is similar to the Omicron BA.1 spike trimer, despite about 22 different mutations between the spike protein of BA.2 and BA.1. As we would expect, JMB2002 effectively blocked the entry of the Omicron BA.2 pseudovirus into human ACE2-expressing cells in a pseudovirus neutralization assay, with the half-maximal inhibition concentration (IC_50_) of 0.2 μg/ml (Fig. 3b), which is the same as for the inhibition of Omicron BA.1 pseudovirus. Taken together, JMB2002 is a broad-spectrum anti-SARS-CoV-2 antibody that has equal inhibition efficacy to both Omicron BA.1 and BA.2 strains.

**FIG 3.**
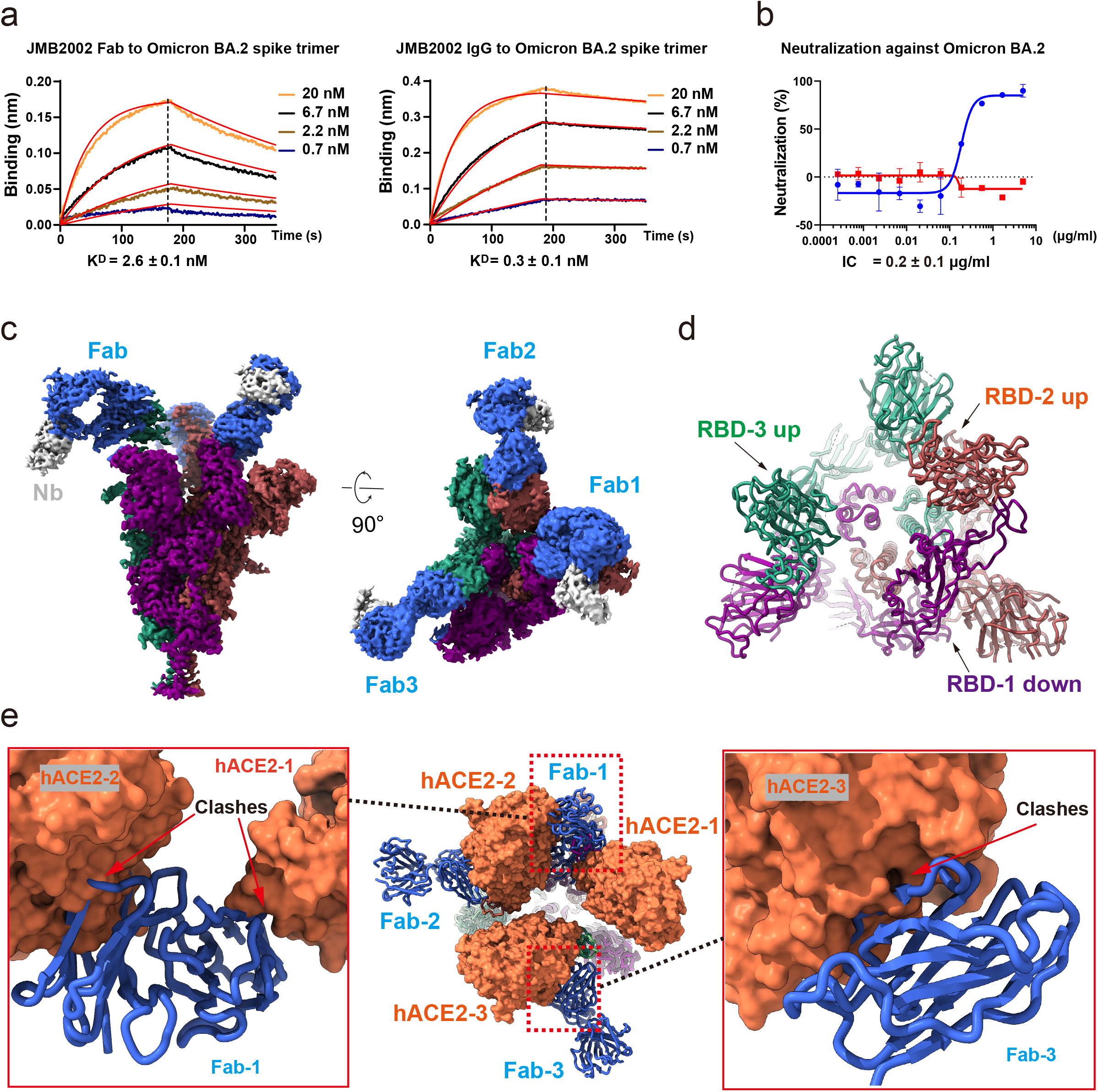
Inhibition of ACE2 binding to the Omicron BA.2 spike trimer by the anti-Omicron antibody JMB2002. **a**. Binding of JMB2002 Fab and IgG to the Omicron and WT spike trimer. **b**. Inhibition of the pseudovirus of Omicron BA.2 by JMB2002. **c**. Cryo-EM density map of the Fab-bound Omicron BA.2 spike trimer shown as front and top views. **d**. Top view of Fab-bound Omicron BA.2 spike trimer complex model with Fab and nanobody hidden. **e**. Superposition of the ACE2-bound and Fab-bound Omicron BA.2 spike trimer showing that Fab binding to RBD inhibits ACE2 binding

To explore the basis of JMB2002 inhibition of Omicron BA.2, we solved the structure of the Omicron BA.2 spike trimer bound to a Fab from JMB2002 at a global resolution of 3.27 Å (Fig. 3c, and Supplementary information, Fig. S5), with the aid of the same nanobody to stabilize the constant regions of Fab as we used in the Omicron BA.1 study.^3^ The cryo-EM density map reveals the binding of three Fab molecules, with each RBD (two RBDs up and one RBD down) bound to a Fab (labeled as RBD-1-3 and Fab-1-3 in Figs. 3c and 3d), which was also similar to one of our previously solved structures of JMB2002 Fab bound to the Omicron BA.1 spike trimer.^3^ Superposition of the structure of Fab-bound Omicron BA.2 spike trimer with the 3-hACE2-bound BA.2 structure demonstrated that Fab-1 binding to the down RBD-1 would clash with hACE2-1 and hACE2-2, while Fab-3 binding to the up RBD-3 would clash with hACE2-3 (Fig. 3e). Thus, the binding of three Fabs to the spike trimer would completely block ACE2 binding.

### Cross-species binding of ACE2 with the Omicron spike trimer

Omicron variants have been proposed to evolve independently from previous VOCs and their origins are elusive.^3^ To investigate the potential evolution roadmap of Omicron variants, we evaluated the cross-species ACE2 binding to the spike trimer from WT and Omicron BA.1 and BA.2 strains. Unlike human ACE2, which showed enhanced binding to the Omicron spike trimer, ACE2 from horse, pig, and sheep displayed decreased binding, with cat ACE2 protein showing similar binding between WT and Omicron variants (Supplementary information, Fig. S2). Meanwhile, both the rat and dog ACE2 showed no binding with WT, Omicron BA.1, or Omicron BA.2 trimers. To our surprise, we found that mouse ACE2 bound to both Omicron BA.1 and BA.2 spike trimer with high affinity (Figs. 4a and 4b), but no binding to WT spike trimer (Fig. 4c). Moreover, mouse ACE2 binds to the Omicron BA.2 spike trimer with approximately threefold increased affinity (*K*_D_ = 2.9 ± 0.2 nM) over the Omicron BA.1 spike trimer (*K*_D_ = 9.1 ± 7.1 nM) (Figs. 4a, 4b and 4d).

**FIG 4.**
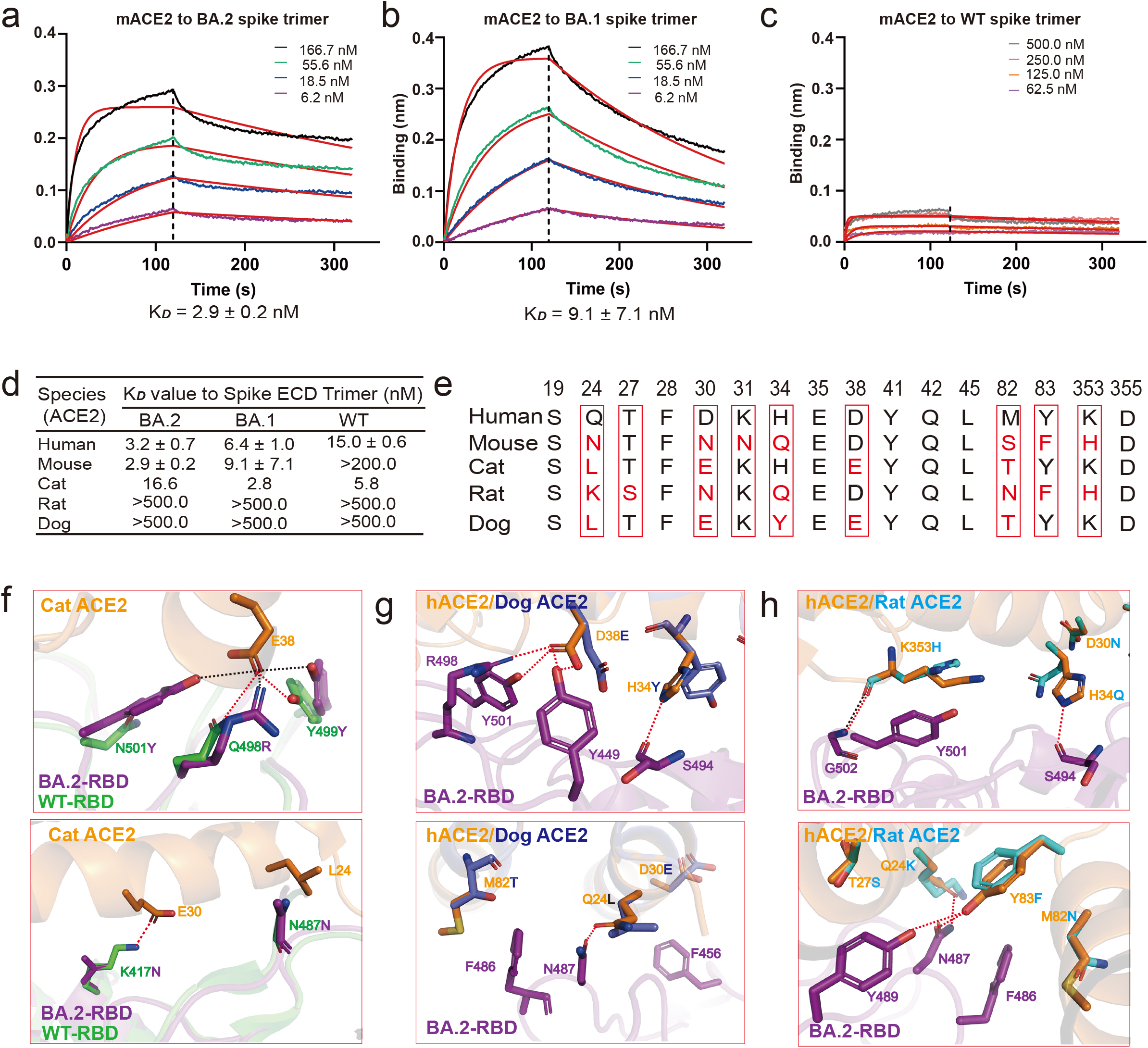
Characterization of the binding affinity between mouse ACE2 and Omicron spike trimers. (**a-c**) Binding of Omicron BA.2 (a), Omicron BA.1(b) and WT (c) spike trimers to mouse ACE2 as determined by BLI. **d**. The *K*_D_ values of tested pairs in this study as determined by BLI. **e**. Amino acid alignment of the 16 key residues in hACE2 with 4 ACE2 orthologs from mouse, cat, rat, and dog. **f**. Superposition of the structures of human ACE2 bound Omicron BA.2 spike trimer and the cat ACE2 bound original spike trimer. (**g and h**) The binding modes of dog (g) and rat (h) ACE2s with the BA.2 RBD, were generated based the complex structure of human ACE2 bound Omicron BA.2 spike trimer. The differentiated residues among ACE2 protein that are responsible for the interactions with SARS-CoV-2 spike protein are showed with detained interactions.

Multiple sequence alignment reveals that 7 out of 16 key interface residues of ACE2 residues involved in interactions with the SARS-CoV-2 spike trimer are highly conserved in human, mouse, cat, rat, and dog, which include S19, F28, E35, Y41, Q42, L45, and D355 (Fig. 4e and Supplementary information, Fig. S2). The rest 9 residues are varied from species to species, thus leading to a diverse binding behavior to the SARS-CoV-2 spike trimer. A homology model of cat ACE2 bound to the BA.2 RBD, based on the solved BA.2 RBD in this work and cat ACE2 structure (Fig. 4f),^29^ revealed that cat ACE2 can make the similar hydrogen bonds with RBDs from WT and both Omicron strains, resulting in comparable binding activity. However, homologous ACE2 residues from dog (L24, Y34, and E38) and rat (K24, Q34, and F83) cannot make similar interactions to the BA.2 RBD, leading to no binding to the BA.2 spike trimer, consistent with our binding data (Figs. 4d, 4e, 4g and 4h and Supplementary information, Fig. S2). In aggregate, our results show the determinant residues of ACE2 for cross-species specificity.

### Cryo-EM Structures of BA.2 and BA.1 spike trimer binding with mACE2

To reveal the molecular basis for the high affinity binding of the Omicron BA.2 and BA.1 spike trimer to mouse ACE2, we determined their complex structures (Figs. 5a-5d, Supplementary information, Figs. S6 and S7). Two major states were observed for both BA.2-mACE2 and BA.1-mACE2 complexes, in which the spike trimer binds to one or two mACE2. In the first state, one RBD is in the open up position with mACE2 binding, and the other two RBDs are in the close down conformation without mACE2 binding (Figs. 5a and 5c). The second state is with two RBDs in the open up position with mACE2 binding, and the third RBD is in the close down position without mACE2 binding (Figs. 5b and 5d). We observed similar RBD-RBD interactions in all mACE2 complexes with the spike trimer of BA.1 and BA.2, which is consistent with the similar interaction in Omicron spike trimer-hACE2 complexes. A BA.2-mACE2 complex structure with all three RBDs bound to mACE2 in the open up conformation was also obtained at 4.5 Å resolution upon further particle classification (Supplementary information, Fig. S6e). Local refinement of the RBD-ACE2 region produced a high-quality map of BA.2 RBD-mACE2 and BA.1 RBD-mACE2 at 2.37 Å and 3.01 Å resolution, respectively, which allowed unambiguous model buildings of the RBD-mACE2 complexes (Figs. 5e, 5f, Supplementary information, Figs. S6 and S7). Structure comparison of the BA.2 RBD-mACE2 complex with the BA.1 RBD-mACE2 complex shows a similar overall organization, which also resembles the Omicron RBD-hACE2 complexes.

**Fig 5.**
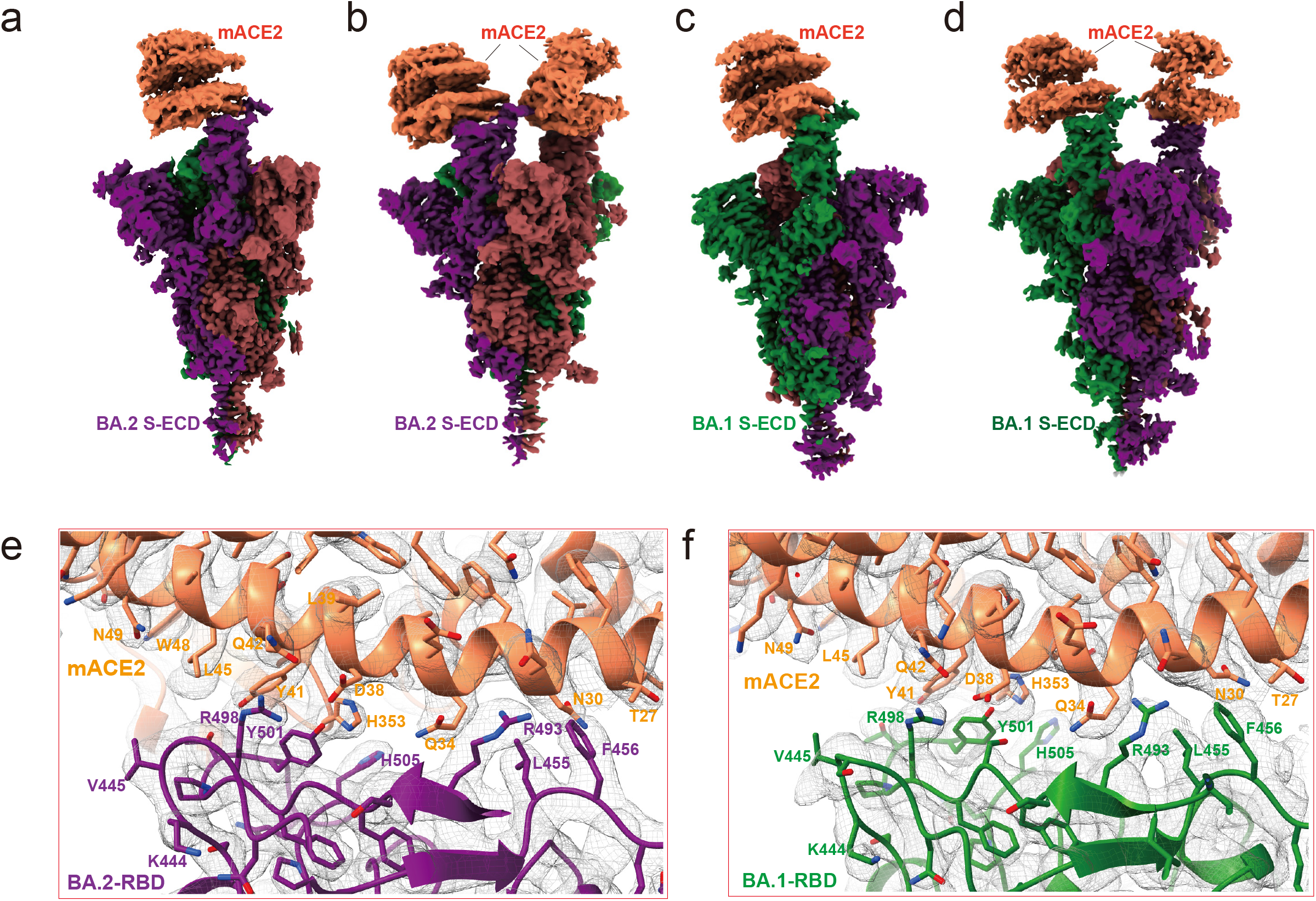
Cryo-EM structure of the Omicron BA.2 and BA.1 spike trimers in complex with mACE2. (**a and b**) Cryo-EM maps of the Omicron BA.2 spike protein-mACE2 complex with one RBD in “up” conformation at 3.2-Å resolution (a), and mACE2 complex with two RBDs in “up” conformation at 3.3-Å resolution (b), respectively. The three protomers are colored in purple, red and green, the density for mACE2 is colored in coral. (**c and d**) Cryo-EM maps of the Omicron BA.1 spike protein-mACE2 complex with one RBD in “up” conformation at 3.1-Å resolution (c), and mACE2 complex with two RBD in “up” conformation at 3.2-Å resolution (d), respectively. The three protomers are colored in purple, red, and green, the density for mACE2 is colored in coral. (**e and f**) Density maps and atomic models of the interaction interface in the BA.2 spike trimer-mACE2 and BA.1-mACE2 complexes.

### Molecular interactions between mouse ACE2 and Omicron RBDs

The high affinity binding of mACE2 to both BA.2 and BA.1 spike trimers can be rationalized with the extensive interactions between mACE2 and the RBDs from both Omicron strains. In the BA.2 RBD-mACE2 structure, Y449 of BA.2 RBD forms two hydrogen bonds with D38 and Q42 of mACE2, N487 of BA.2 RBD forms a hydrogen bond with N24 of mACE2, while T500 of BA.2 RBD forms two hydrogen bonds with Y41 and D355 of mACE2 (Figs. 6a and 6b). Meanwhile, the BA.2 RBD forms additional interactions with mACE2 from the mutated residues. Particularly, Q493R forms three hydrogen bonds with N31 and Q34, and a salt bridge with E35 of mACE2; Q498R forms a salt bridge with D38, and a hydrogen bond with Q42 of mACE2; while N501Y forms extensive π-π stacking interactions with H353 of mACE2 (Figs. 6a and 6b). The interactions of BA.1 RBD with mACE2 closely resemble those in the BA.2 RBD-mACE2 complex (Figs. 6c and 6d). The interactions of the RBDs from both Omicron strains with mACE2 cannot be completely satisfied in the hypothetical model of the WT RBD-mACE2 complex (Figs. 6e and 6f), which are consistent with our binding assays that the WT RBD cannot bind to mACE2. Collectively, our results indicate that the three mutations of Q493R, Q498R, and N501Y in the Omicron RBDs are essential for mACE2 binding.

**Figure 6.**
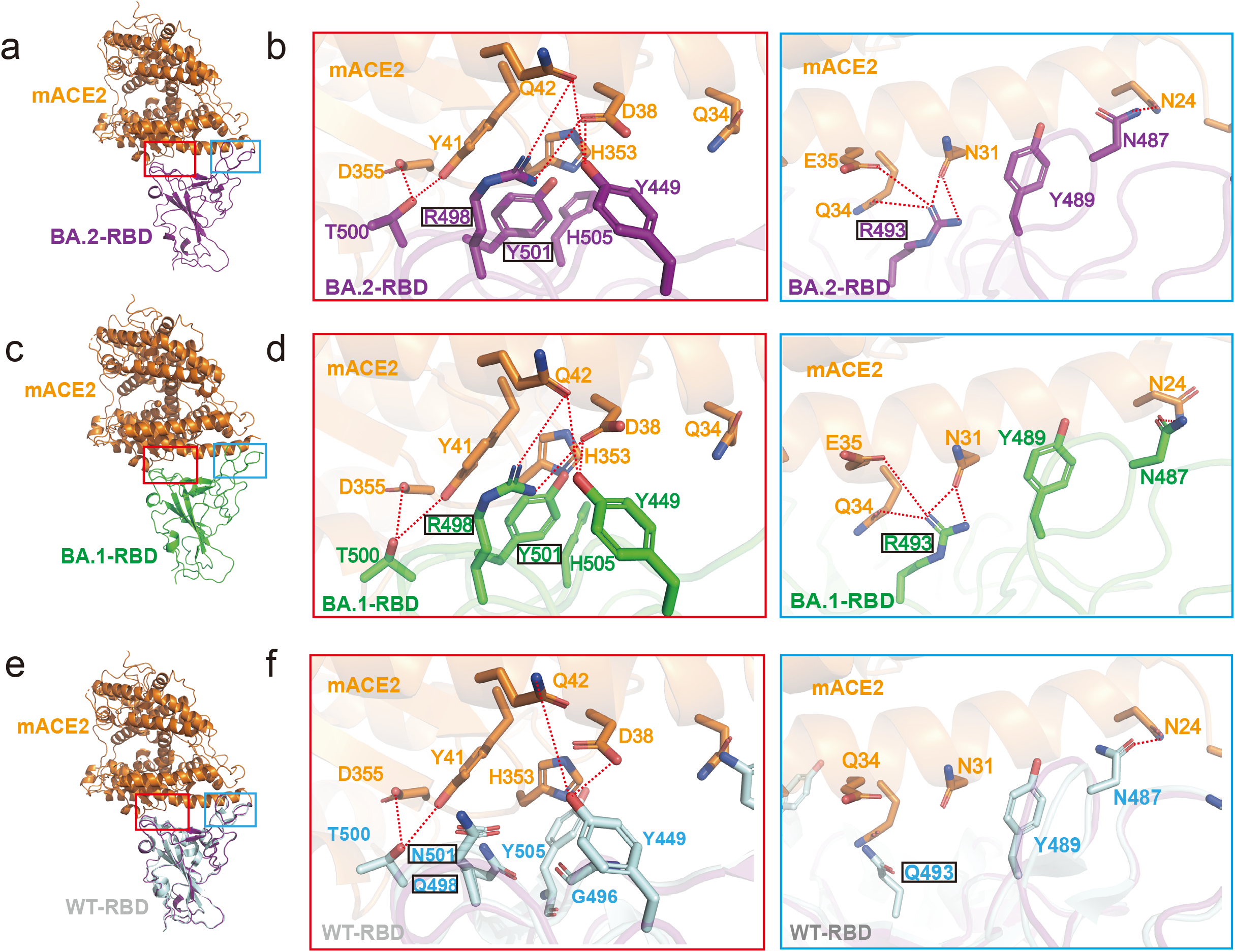
Structural analysis of mACE2 and RBD. **a**. Overall structure of the BA.2-RBD and mACE2 complex. **b**. Details of the binding between BA.2-RBD and mACE2. The binding between BA.2-RBM and mACE2 consists mainly of two interaction regions, marked in a red or a blue box. Residues involved in the interactions are shown as sticks. Hydrogen bonds are shown as dashed black lines. (**c and d**) Overall structure of the BA.1-RBD and mACE2 complex, with detailed hydrogen bond or salt bridge interactions in BA.1-RBD and ACE2 interface with the same view as in (a and b). (**e and f**) The superposition of the BA.2 RBD and mACE2 complex with SARS-CoV-2 WT-RBD alone (PDB: 6LZG). Detailed hydrogen bond or salt bridge interactions are highlighted with the same view as in (a and b).

Given that ACE2 is highly conserved among mammals including mouse and human, we aligned the two structures of BA.2 RBD-hACE2 and BA.2 RBD-mACE2 with the root mean square deviation of 0.48 Å for 670 Cα atoms from both RBD and ACE2. Detailed structural analysis reveals a similar interaction network from mACE2 and hACE2 to the BA.2 RBD (Figs. 7a and 7b), supporting that both mACE2 and hACE2 bind to the BA.2 RBD with high affinity.

**Figure 7.**
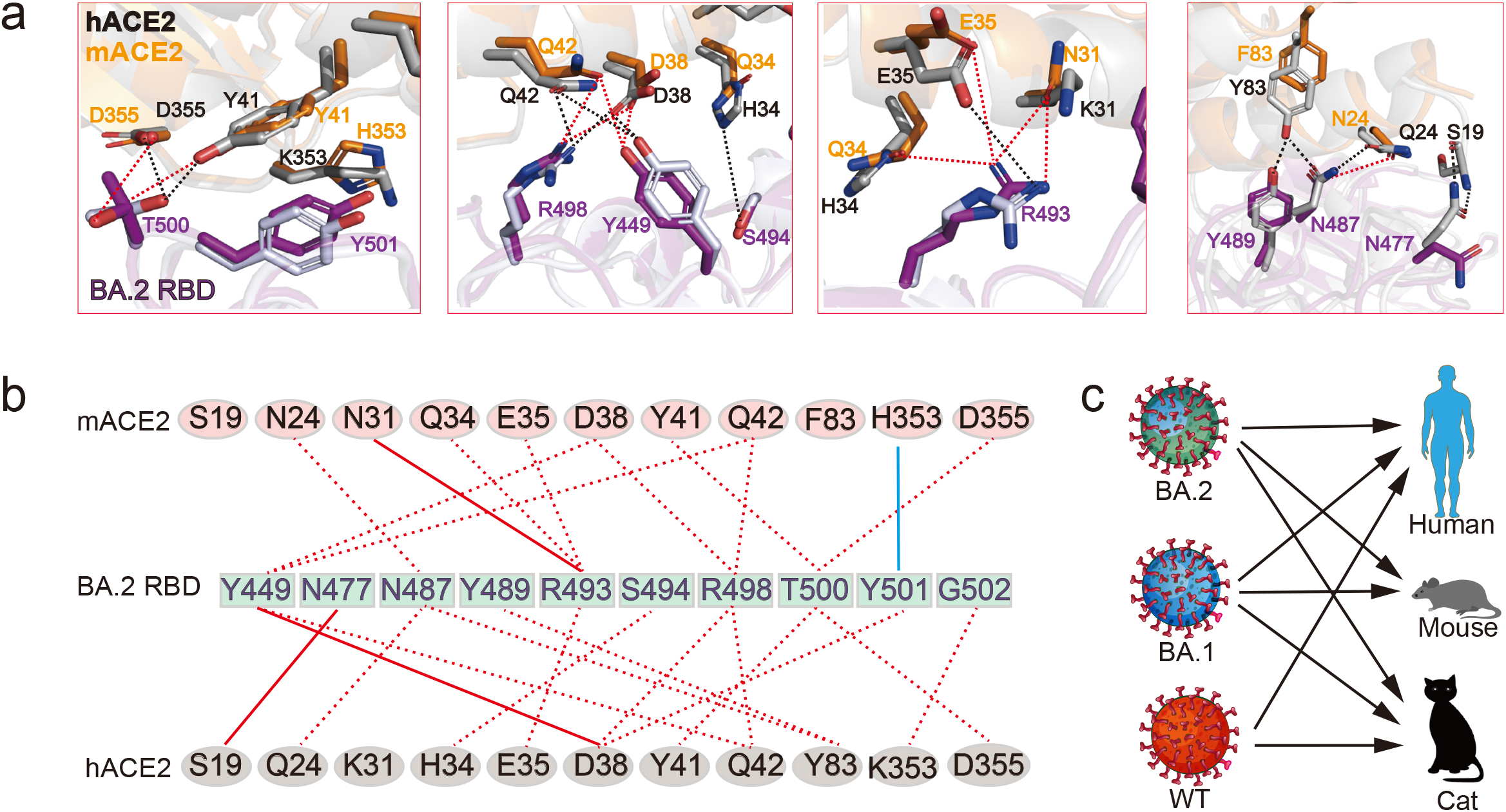
Comparison of the binding mode of the BA.2 RBD to mACE2 and hACE2. **a**. The superposition of BA.2 RBD bound to mACE2 with BA.2 RBD bound to hACE2, with hydrogen bond or salt bridge interactions shown as dashed lines. Hydrogen bond or salt bridge interactions in hACE2-RBD and mACE2-RBD are shown as red and black dashed lines, respectively. **b**. Residues involved in the interactions of BA.2 RBD with hACE2 and mACE2 are listed and connected by lines. Red dashed lines indicate one hydrogen bond or salt bridge and red solid lines indicate two hydrogen bonds, whereas the cyan solid line represents a π-π stacking interaction between the residues. **c**. Infection selectivity of BA.1, BA.2 and WT strains to human, mouse and cat.

## DISCUSSION

BA.2, a new variant of Omicron, has rapidly spread throughout the world and has become the dominant strain due to its higher infectivity than BA.1. In this study, we first biochemically analyzed the binding affinity of the BA.2 spike trimer to hACE2, which showed around 2-fold higher affinity than that of BA.1 and 11-fold higher affinity than that of the WT strain. The spike-ACE2 interaction is the first step of viral binding to host cells, which is critically important for viral entry and subsequent infections. Thus, the higher binding potency of the BA.2 spike trimer likely contributes to its higher transmission capability although many other factors such as spike trimer could also contribute to viral entry and infectivity. The structure of the BA.2 spike trimer with hACE2 reveals an extensive interaction network in the RBD-hACE2 interface, which is also conserved in the BA.1 spike trimer-hACE2 complex. The network of interaction of the BA.2 spike trimer with hACE2 is more extensive than the WT spike trimer-ACE2 complex, providing a structural basis for the high binding potency of the BA.2 spike trimer to hACE2. In addition, the RBD of BA.2 is more stable than the BA.1 RBD because of an additional interaction of R403 with D405N in BA.2, which is missing in BA.1 (Fig. 1e). The higher stability of the BA.2 RBD than that of the BA.1 RBD might also contribute to the higher binding affinity of the BA.2 RBD to hACE2 because the high stability of the structure reduces the binding energy cost associated with entropy loss upon binding.

Like the BA.1 strain, BA.2 also harbors many mutations that help its decreased sensitivity to many neutralizing monoclonal antibodies (mAbs). In addition, 4 missense mutations of S371F, T376A, D405N, and R408S in BA.2 RBD but not in BA.1 RBD could also potentially increase the immune evasion of neutralizing antibodies and vaccination, which brings even more challenges to effectively subdue the current COVID-19 pandemic. On the other hand, the spike protein of both BA.1 and BA.2 has 37 and 31 mutations out of 1273 residues, representing less than 3% of total mass. Although most Omicron mutations are clustered on the epitopes to many neutralizing monoclonal antibodies, the majority of the spike protein surface resembles the WT spike protein (Figs. 1f and 1g). Using the WT spike protein as the antigen for vaccines can induce B cell and T cell responses,^30^ which could recognize the overall structures of the spike trimer as well as other conserved regions outside of the hot epitope regions. Although vaccination with the WT spike protein as antigen as used in many current vaccines cannot completely block Omicron infections, it can reduce severity of Omicron infection, possibly through the remaining immunity induced by the WT spike protein.

Despite less severe symptoms in the general population infected with Omicron variants, immunocompromised people with underlying diseases are still at high risk of developing severe forms of COVID-19. Antibody therapy could serve as a viable option for these patients either as a preventive treatment or a therapeutic option. However, several monoclonal antibodies available in clinical practice have shown decreased potency to Omicron BA.1 and BA.2 variants.^31^ Thus, broad-spectrum anti-SARS-CoV-2 antibodies, especially with potent activities against Omicron variants are currently in urgent need. Strikingly, we reconfirmed the strong efficacy of our previously discovered antibody, JMB2002, against Omicron BA.2 in both binding assays and pseudovirus neutralization assay, with comparable inhibition potency to Omicron BA.1 strain. Further structural studies of JMB2002 Fab-bound Omicron BA.2 spike trimer enabled us to propose the inhibition mechanism of JMB2002 against Omicron BA.2, which is similar to the inhibition of Omicron BA.1 by JMB2002. As for Omicron BA.1, structural analysis reveals that the binding of JMB2002 (IgG or Fab) to the BA.2 spike trimer can completely block ACE2 binding, thus providing the basis for inhibition of both Omicron BA.1 and BA.2 variants by JMB2002.

In addition, our results also provided a possible origin host for Omicron BA.1 and BA.2. As reported, Omicron variants evolved independently of all other VOCs and its origin is a mystery.^3^ Investigation of the origin of Omicron BA.1 and BA.2 is critical for the effective control and prevention of COVID-19 in the human population. Mutation profile analysis suggests that the Omicron variants may have arisen through evolution via the mouse as a host.^4^ In this study, our biochemical data confirmed that both BA.1 and BA.2 spike trimers are able to bind with high affinity to mouse ACE2 and cat ACE2, while the original WT spike binds well to cat ACE2 but not to mouse ACE2, indicating high susceptibility of mouse and cat to BA.1 and BA.2 infections (Fig.7c). We determined the structures of spike trimers from both BA.1 and BA.2 in complex with mouse ACE2, and our structures elucidated that the three residue mutations Q493R, Q498R, and N501Y are essential for mACE2 binding, suggesting their importance in infections. Intriguingly, mutations on these three residues Q493, Q498, and N501, were detected in the mouse-adapted SARS-CoV-2.^32-34^ Surprisingly, the Q498R and Q493R mutations were detected in mouse-adapted SARS-CoV-2 by passage 10 and passage 20, respectively, and have not since been detected in any SARS-CoV-2 variants from other animals.^35^ Based on our data reported here, combined with previous studies, we propose mouse is likely a host for evolution of Omicron variants.

Taken together, our data reveal structural and biochemical insights into the enhanced transmissibility and antibody inhibition of Omicron BA.2 as well as a possible evolutionary pathway for Omicron variants. We propose that Omicron variants may pass from human to cat, then to mouse, and then back to human (Fig.7c). Although such an evolution pathway is highly speculative, the ability for Omicron variants to infect and spread in mice and possibly other animals could have important implications in the establishment of control strategy in combating SARS-CoV-2 infection.

## Supporting information

combined file of the full-length paper

## AUTHOR CONTRIBUTIONS

Y.X., C.W. and W.Y. designed the expression constructs, purified the spike complexes, screened the cryo-EM conditions, prepared the cryo-EM grids, and participated in figure and manuscript preparation. Y.X. collected cryo-EM images with the help of Q.Y., K.W., Y.X. performed density map calculations, built and refined the models. H.L. participated in figure and manuscript preparation. C.W., M.J. and X.X.W. purified the ACE2. S-J. D. supervised X.C., C.G., J.L., D.W., X.H. and X.P.W., provided the JMB2002 antibodies, performed the function assays for JMB2002 antibodies and spike trimers, and participated in manuscript writing; H.E.X. and W.Y. conceived and supervised the project, analyzed the structures, and wrote the manuscript with inputs from all authors.

## ACKNOWLEDGMENTS

The cryo-EM data of the Omicron spike protein complexes were collected at the Shanghai Advanced Electron Microscope Center, Shanghai Institute of Material Medica. This work was partially supported by Ministry of Science and Technology (China) grants (2018YFA0507002 to H.E.X.); Shanghai Municipal Science and Technology Major Project (2019SHZDZX02 to H.E.X.); Shanghai Municipal Science and Technology Major Project (H.E.X.); CAS Strategic Priority Research Program (XDB37030103 to H.E.X.), National Natural Science Foundation of China (32130022 to H.E.X.); the Youth Innovation Promotion Association of CAS (2021278 to W.Y.); National Natural Science Foundation of China (32171189 to W.Y.); National Science Fund for Excellent Young Scholars (82122067 to W.Y.); Key tasks of the Lingang Laboratory (LG202103-03-05 to W.Y.); Key tasks of Lingang Laboratory (LG202101-01-03 to Y. X.); National Natural Science Foundation of China (81902085 to Y. X.); China Postdoctoral Science Foundation Funded Project (2021M703342 to C.W.); Shanghai Post-doctoral Excellence Program (2021429 to C.W.). In addition, this work was partially supported by High-level new R&D institute (2019B090904008), and High-level Innovative Research Institute (2021B0909050003) from Department of Science and Technology of Guangdong Province.

## Data Resources

Materials are available from the corresponding authors upon reasonable request. Density maps and structure coordinates have been deposited in the Electron Microscopy Data Bank (EMDB) and the Protein Data Bank (PDB) with accession codes EMD-XXXX and XXXX for; EMD-XXXX and XXXX for; EMD-XXXX and XXXX for; and EMD-XXXX and XXXX for. Source data are provided with this paper.

