## Supplementary material for "Structural and biochemical mechanism for increased infectivity and immune evasion of Omicron BA.1 and BA.2 variants and their mouse origins": combined file of the full-length paper


#### Competing interests

H.E.X., W.Y., Y.X., C.W., H.L., M.J., X.X.W., Q.Y and K.W. have declared no competing interest. S-J. D., X.C., C.G., J.L., D.W., X.H. and X.P.W., are employee of Shanghai Jemincare Pharmaceuticals, and are developing JMB2002 as a potential anti-Omicron therapeutic.

**Figure 7. Comparison of the binding mode of the BA.2 RBD to mACE2 and hACE2.**

**a.** The superposition of BA.2 RBD bound to mACE2 with BA.2 RBD bound to hACE2, with hydrogen bond or salt bridge interactions shown as dashed lines. Hydrogen bond or salt bridge interactions in hACE2-RBD and mACE2-RBD are shown as red and black dashed lines, respectively.

**b.** Residues involved in the interactions of BA.2 RBD with hACE2 and mACE2 are listed and connected by lines. Red dashed lines indicate one hydrogen bond or salt bridge and red solid lines indicate two hydrogen bonds, whereas the cyan solid line represents a $\pi$-$\pi$ stacking interaction between the residues.

**c.** Infection selectivity of BA.1, BA.2 and WT strains to human, mouse and cat.

### Figure 1

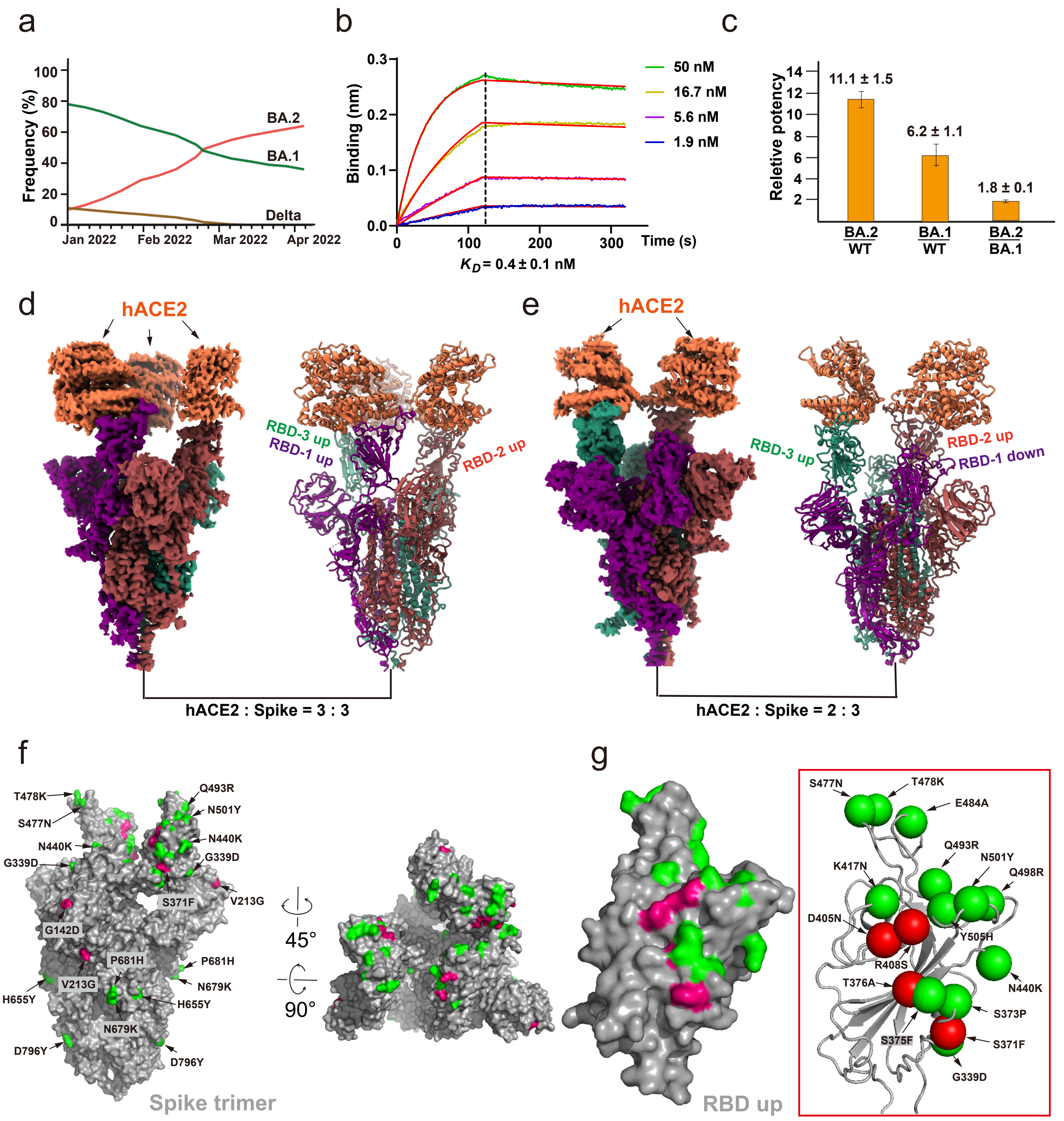

a
Frequency (%)
BA.2
BA.1
Delta
Jan 2022
Feb 2022
Mar 2022
Apr 2022
b
Binding (nm)
50 nM
16.7 nM
5.6 nM
1.9 nM
Time (s)
$K_D$ = 0.4 ± 0.1 nM
c
Reletive potency
11.1 ± 1.5
6.2 ± 1.1
1.8 ± 0.1
BA.2/WT
BA.1/WT
BA.2/BA.1
d
hACE2
RBD-3 up
RBD-1 up
RBD-2 up
hACE2 : Spike = 3 : 3
e
hACE2
RBD-3 up
RBD-2 up
RBD-1 down
hACE2 : Spike = 2 : 3
f
T478K
S477N
N440K
G339D
Q493R
N501Y
S371F
V213G
G142D
P681H
N679K
H655Y
D796Y
45°
90°
Spike trimer
g
RBD up
S477N
T478K
E484A
Q493R
N501Y
Q498R
K417N
D405N
Y505H
R408S
N440K
T376A
S373P
S375F
S371F
G339D

### Figure 2

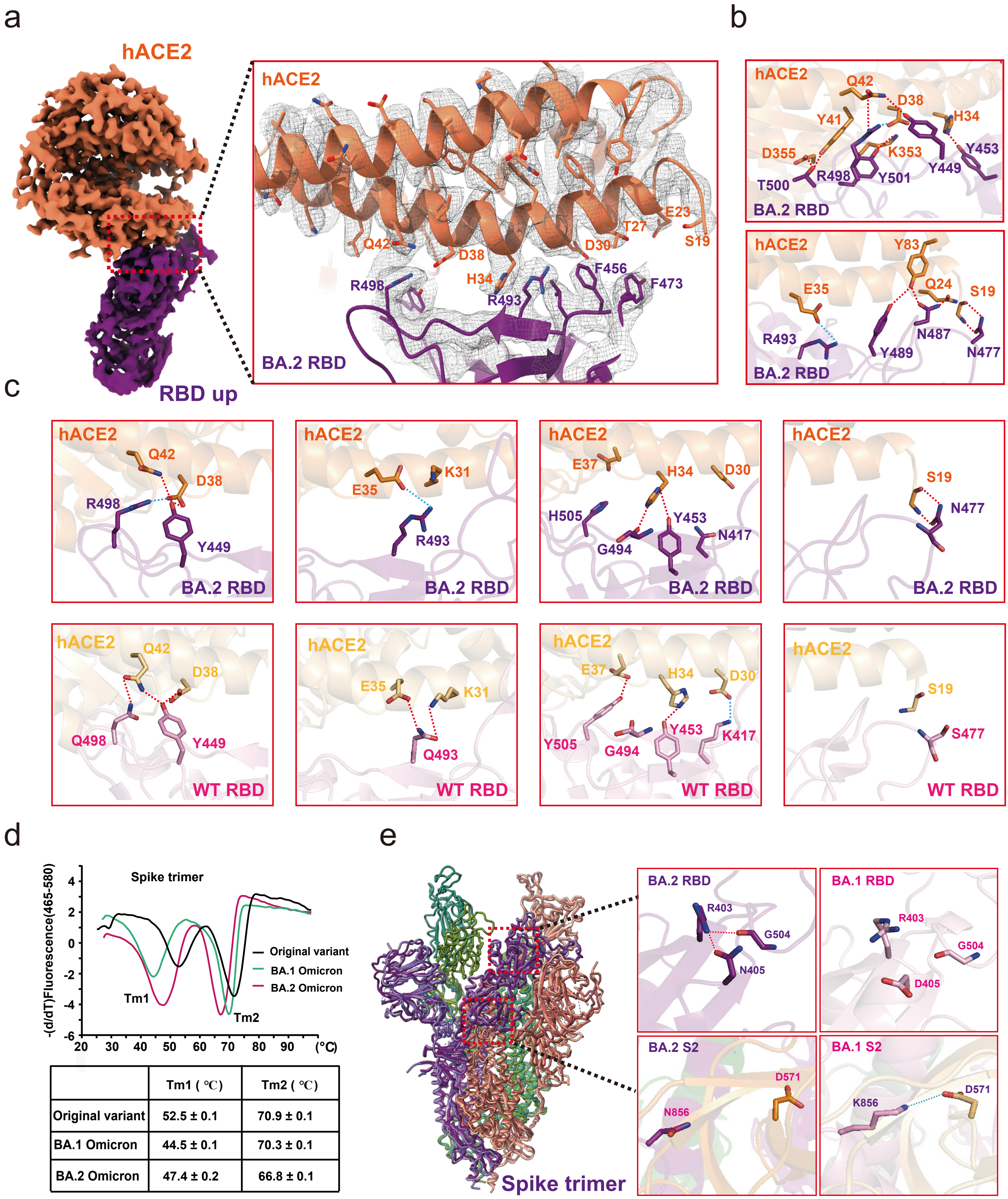


| | Tm1 ( ℃) | Tm2 ( ℃) |
|---|---|---|
| Original variant | 52.5 ± 0.1 | 70.9 ± 0.1 |
| BA.1 Omicron | 44.5 ± 0.1 | 70.3 ± 0.1 |
| BA.2 Omicron | 47.4 ± 0.2 | 66.8 ± 0.1 |

### Figure 3

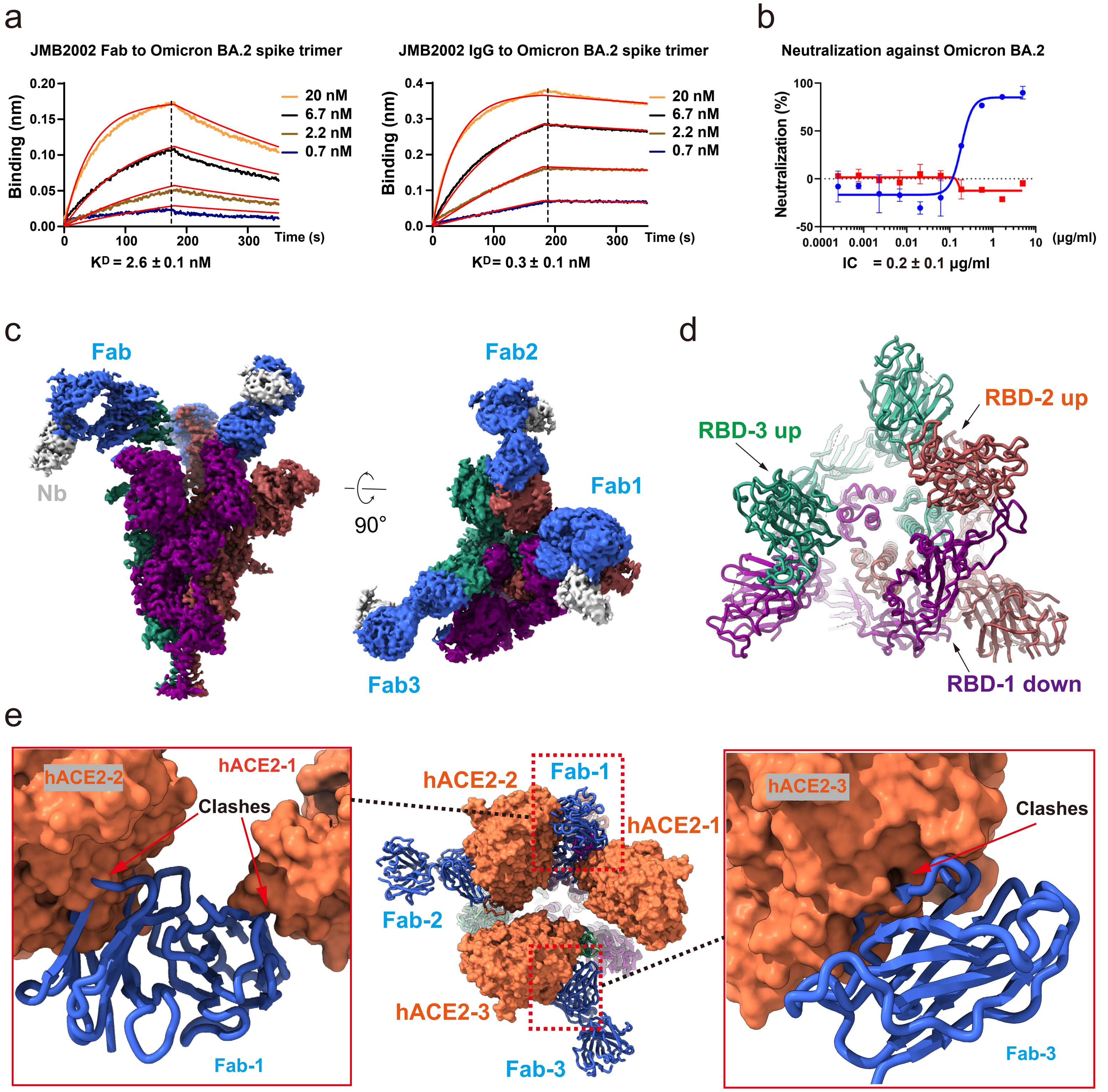
a
JMB2002 Fab to Omicron BA.2 spike trimer
Binding (nm)
20 nM
6.7 nM
2.2 nM
0.7 nM
Time (s)
KD = 2.6 ± 0.1 nM
JMB2002 IgG to Omicron BA.2 spike trimer
Binding (nm)
20 nM
6.7 nM
2.2 nM
0.7 nM
Time (s)
KD= 0.3 ± 0.1 nM
b
Neutralization against Omicron BA.2
Neutralization (%)
(μg/ml)
IC = 0.2 ± 0.1 μg/ml
c
Fab
Nb
90°
Fab2
Fab1
Fab3
d
RBD-2 up
RBD-3 up
RBD-1 down
e
hACE2-2
hACE2-1
Clashes
Fab-1
hACE2-2
Fab-1
hACE2-1
Fab-2
hACE2-3
Fab-3
hACE2-3
Clashes
Fab-3

### Figure 4

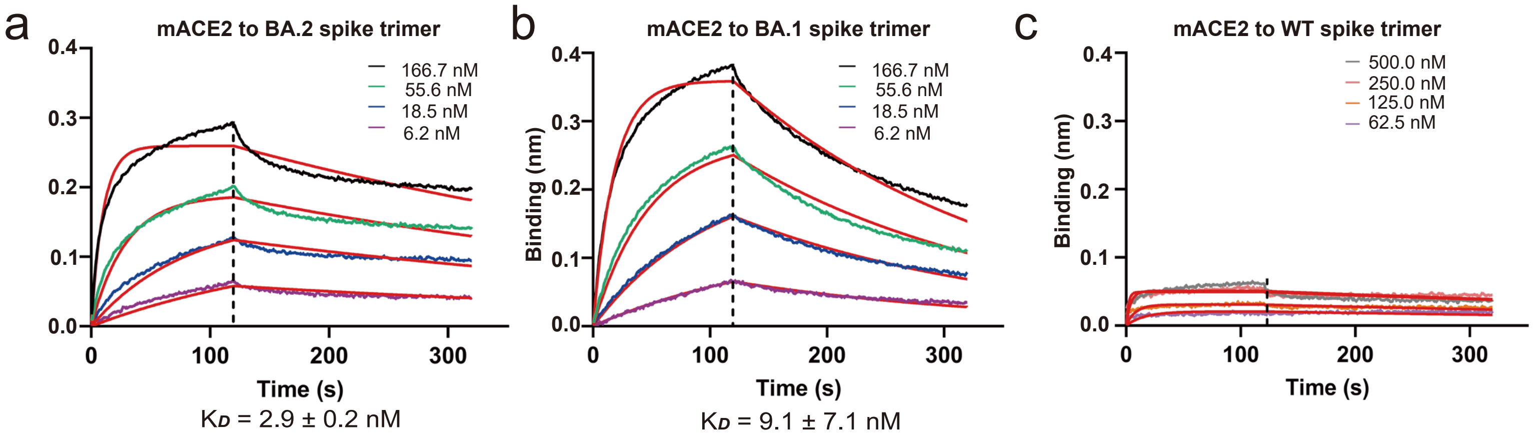


d

| Species (ACE2) | KD value to Spike ECD Trimer (nM) BA.2 | BA.1 | WT |
|---|---|---|---|
| Human | 3.2 ± 0.7 | 6.4 ± 1.0 | 15.0 ± 0.6 |
| Mouse | 2.9 ± 0.2 | 9.1 ± 7.1 | >200.0 |
| Cat | 16.6 | 2.8 | 5.8 |
| Rat | >500.0 | >500.0 | >500.0 |
| Dog | >500.0 | >500.0 | >500.0 |

e

| | 19 | 24 | 27 | 28 | 30 | 31 | 34 | 35 | 38 | 41 | 42 | 45 | 82 | 83 | 353 | 355 |
|---|---|---|---|---|---|---|---|---|---|---|---|---|---|---|---|---|
| Human | S | Q | T | F | D | K | H | E | D | Y | Q | L | M | Y | K | D |
| Mouse | S | N | T | F | N | N | Q | E | D | Y | Q | L | S | F | H | D |
| Cat | S | L | T | F | E | K | H | E | E | Y | Q | L | T | Y | K | D |
| Rat | S | K | S | F | N | K | Q | E | D | Y | Q | L | N | F | H | D |
| Dog | S | L | T | F | E | K | Y | E | E | Y | Q | L | T | Y | K | D |

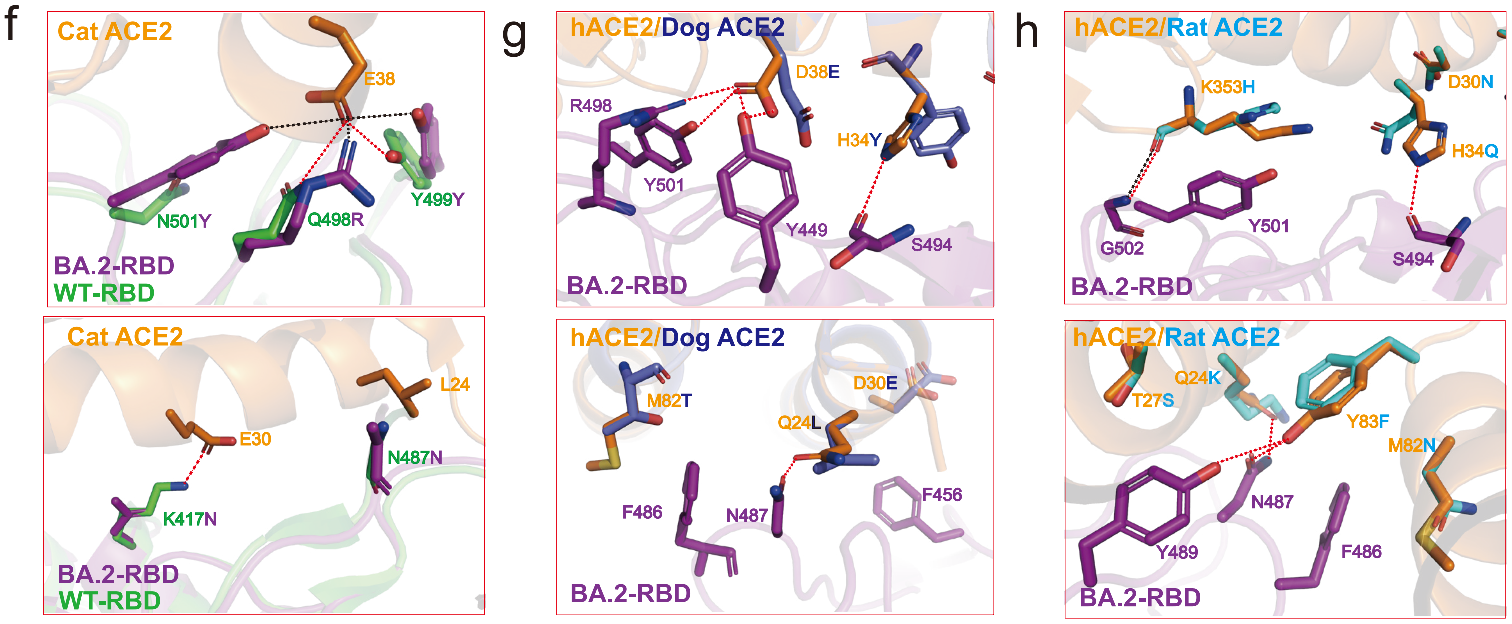

### Figure 5

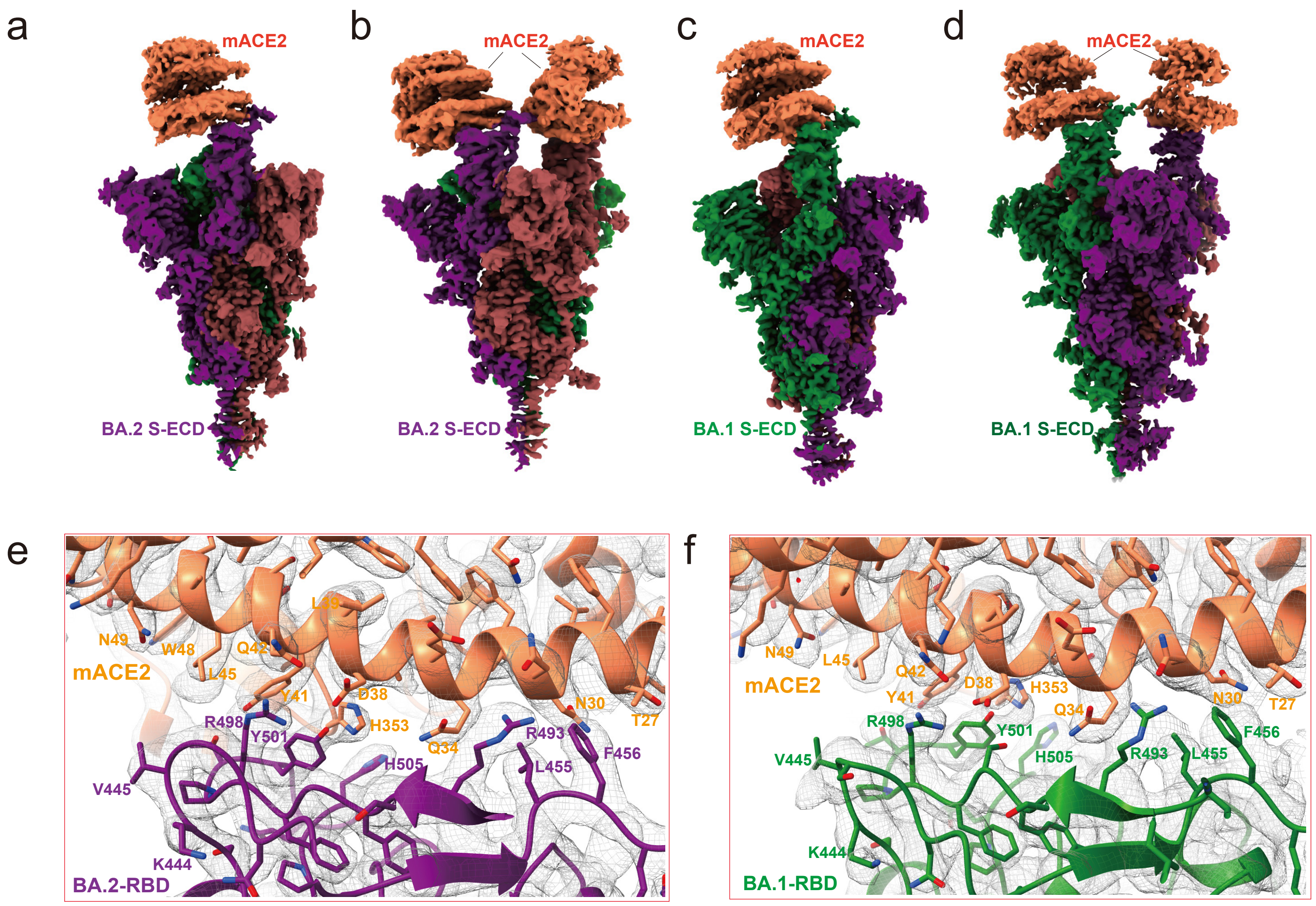
a
mACE2
BA.2 S-ECD
b
mACE2
BA.2 S-ECD
c
mACE2
BA.1 S-ECD
d
mACE2
BA.1 S-ECD
e
mACE2
N49
W48
Q42
L45
L39
Y41
D38
H353
N30
T27
R498
Y501
Q34
R493
F456
H505
L455
V445
K444
BA.2-RBD
f
mACE2
N49
L45
Q42
Y41
D38
H353
Q34
N30
T27
R498
Y501
H505
R493
L455
F456
V445
K444
BA.1-RBD

### Figure 6

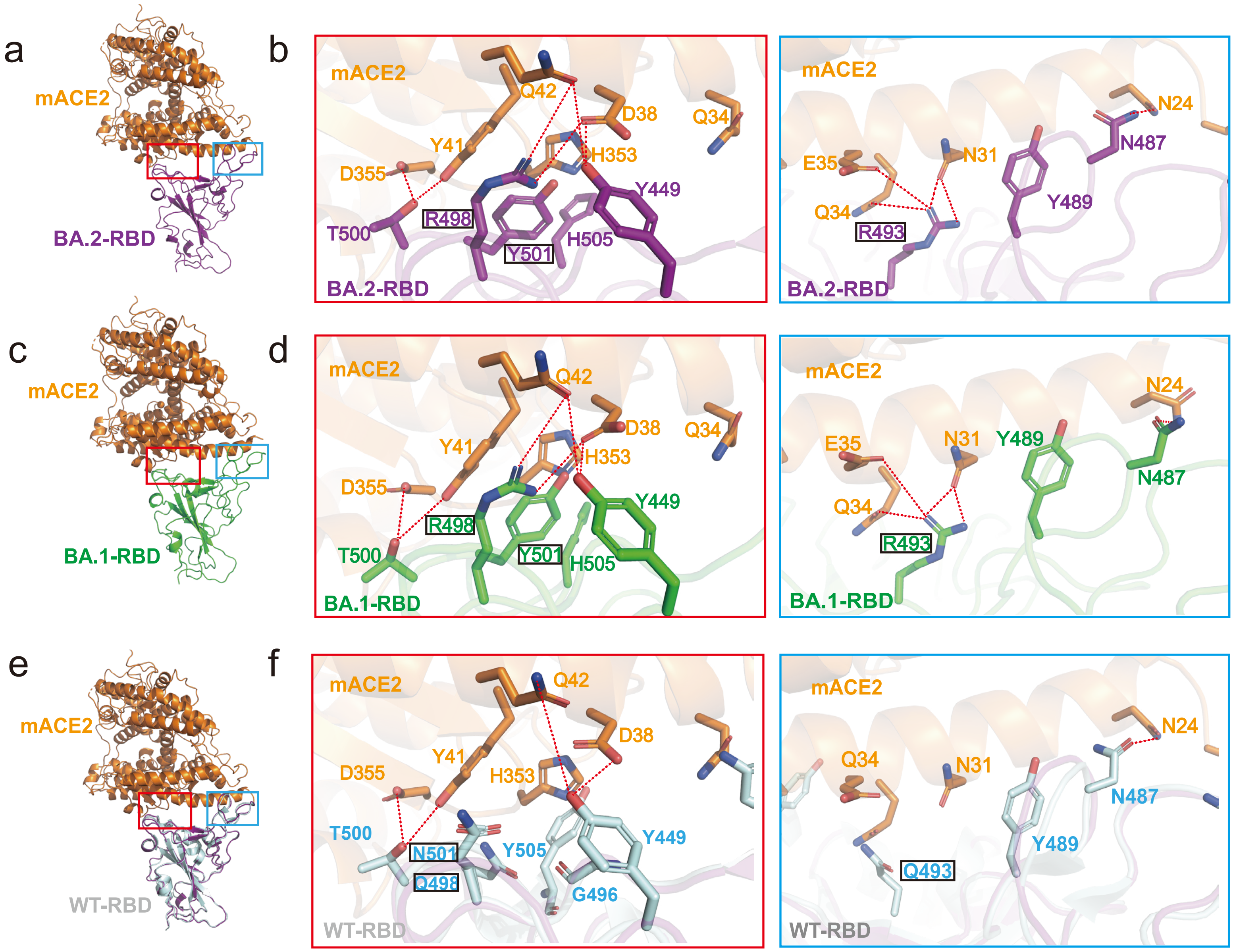
a
mACE2
BA.2-RBD
b
mACE2
Q42
D38
Q34
Y41
H353
D355
Y449
R498
T500
Y501
H505
BA.2-RBD
mACE2
N24
N487
E35
N31
Q34
Y489
R493
BA.2-RBD
c
mACE2
BA.1-RBD
d
mACE2
Q42
D38
Q34
Y41
H353
D355
Y449
R498
T500
Y501
H505
BA.1-RBD
mACE2
N24
E35
N31
Y489
N487
Q34
R493
BA.1-RBD
e
mACE2
WT-RBD
f
mACE2
Q42
D38
D355
Y41
H353
T500
N501
Y505
Y449
Q498
G496
WT-RBD
mACE2
N24
Q34
N31
N487
Y489
Q493
WT-RBD

### Figure 7

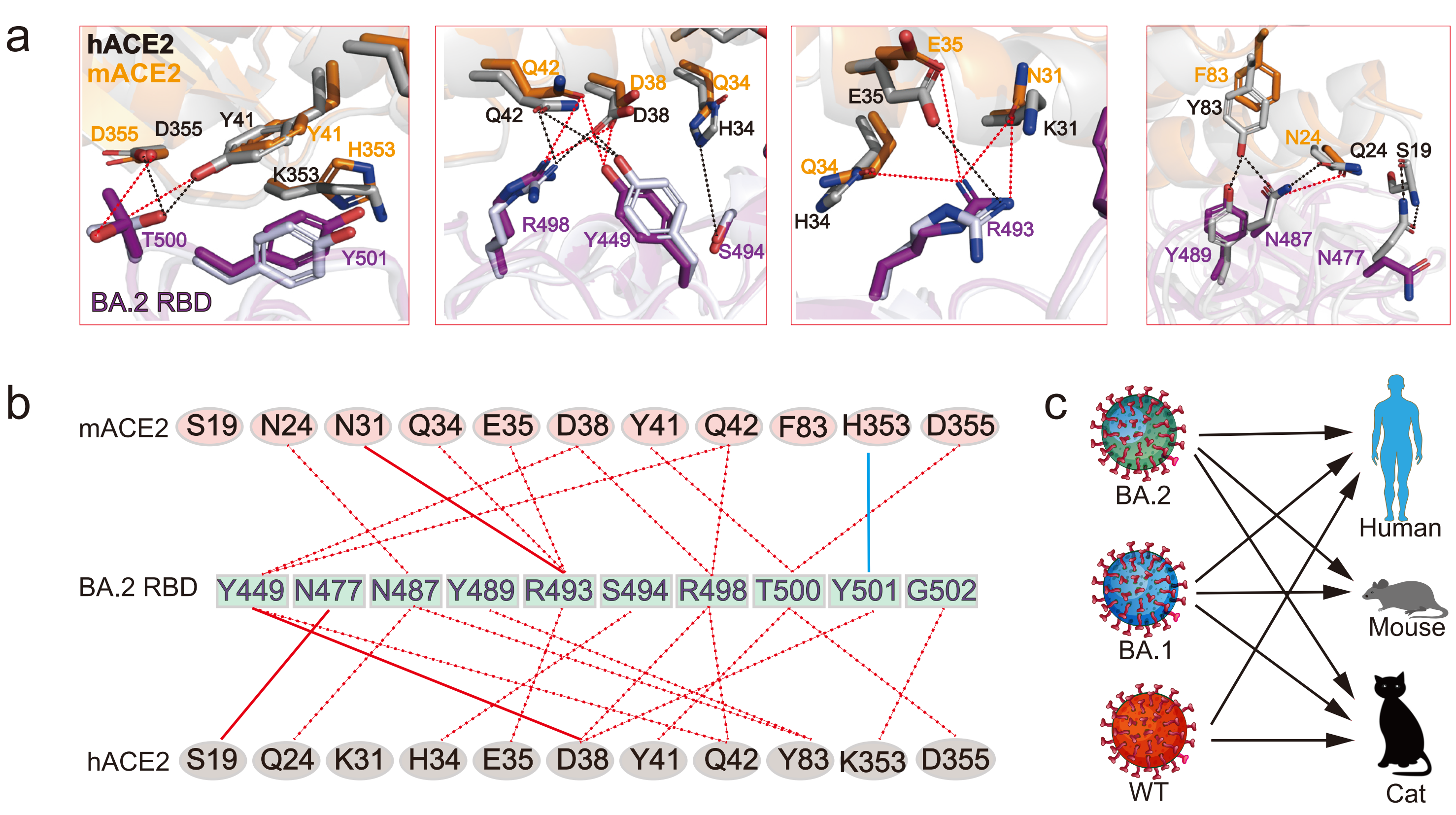

a
hACE2
mACE2
D355
D355
Y41
Y41
H353
K353
T500
Y501
BA.2 RBD
Q42
D38
Q34
Q42
D38
H34
R498
Y449
S494
E35
E35
N31
K31
Q34
H34
R493
F83
Y83
N24
Q24
S19
Y489
N487
N477
b
mACE2
S19
N24
N31
Q34
E35
D38
Y41
Q42
F83
H353
D355
BA.2 RBD
Y449
N477
N487
Y489
R493
S494
R498
T500
Y501
G502
hACE2
S19
Q24
K31
H34
E35
D38
Y41
Q42
Y83
K353
D355
c
BA.2
BA.1
WT
Human
Mouse
Cat

### Supplementary information, Fig. S1

a

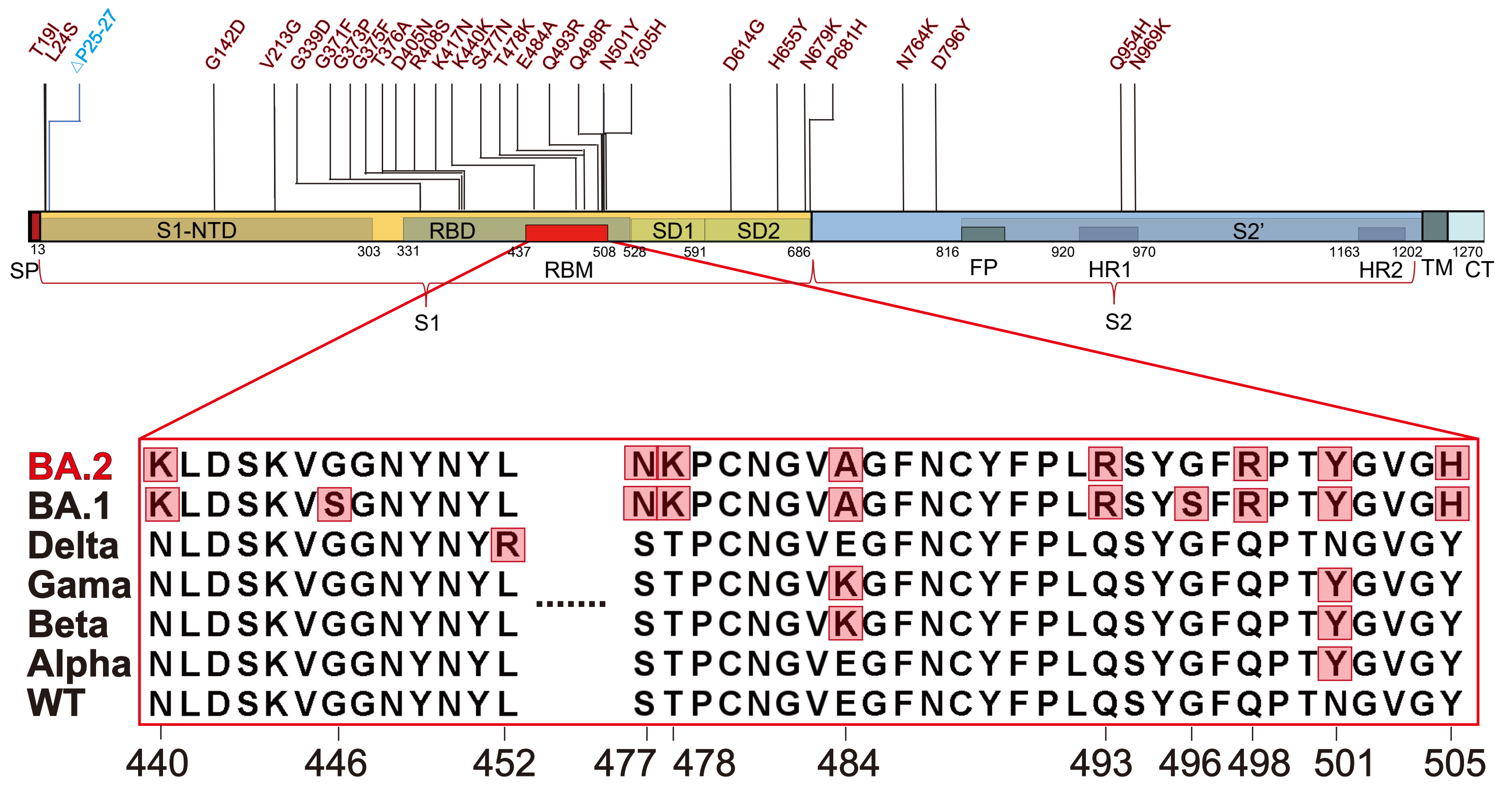

### Supplementary information, Fig. S2

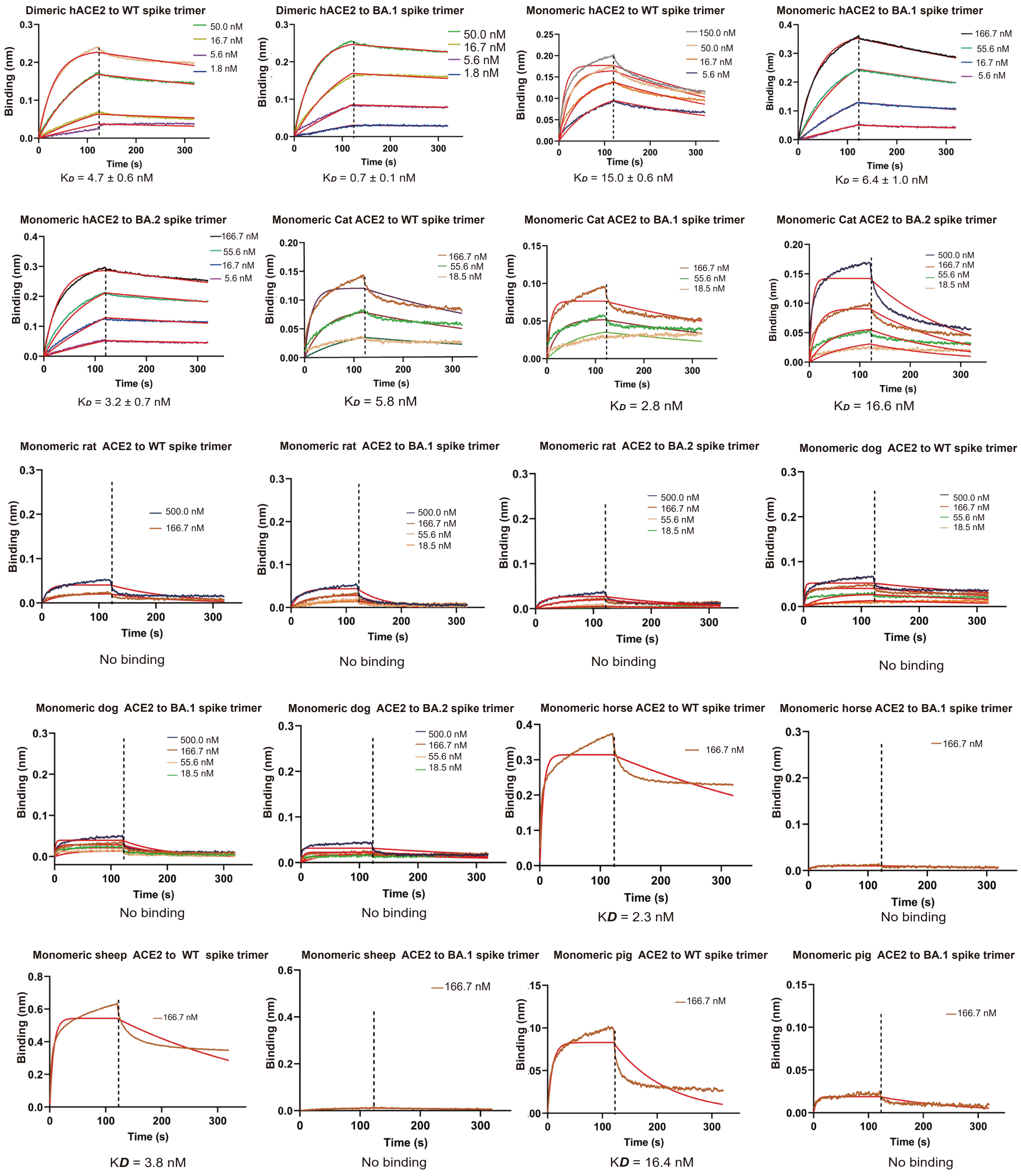

### Supplementary information, Fig. S3

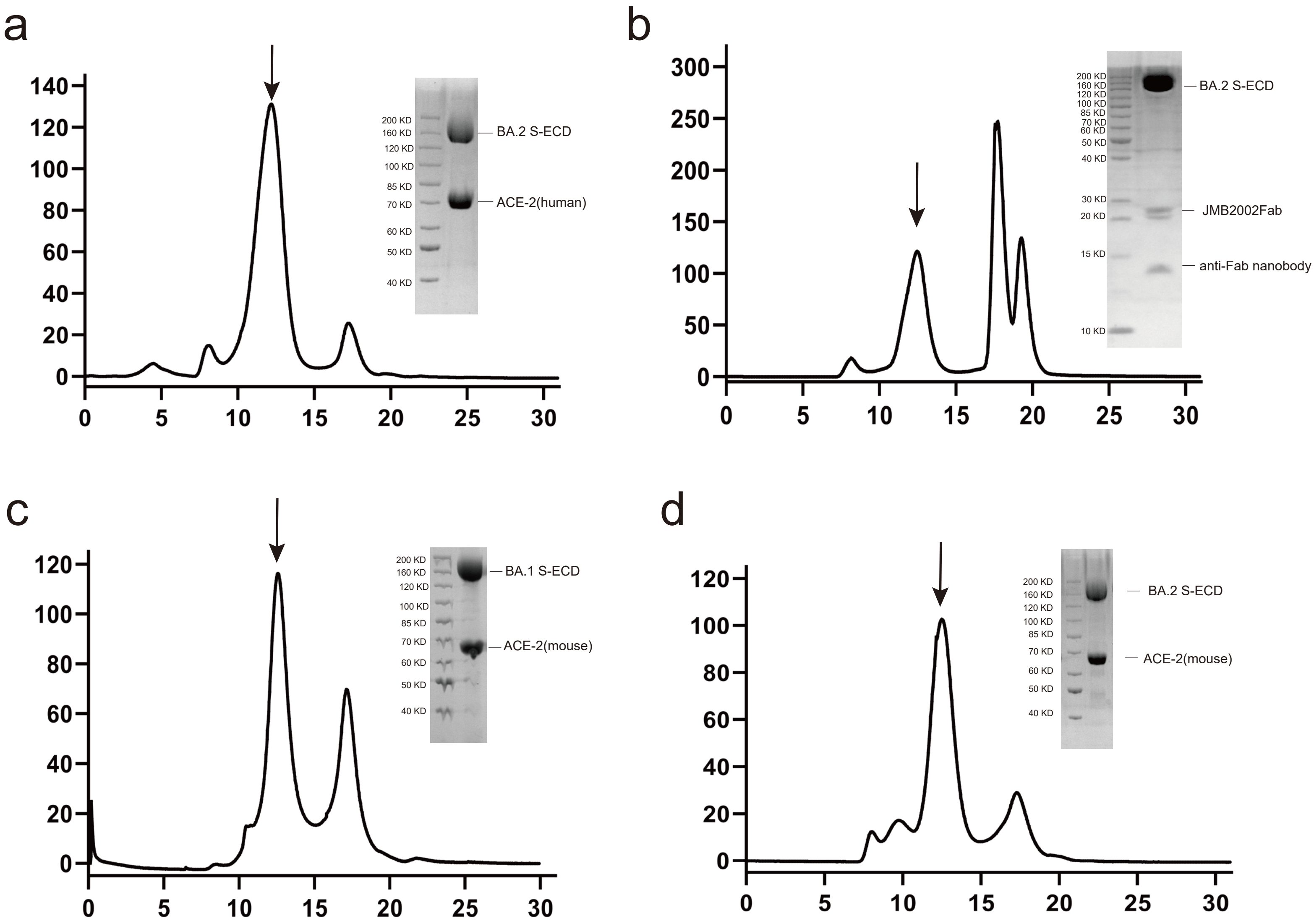

### Supplementary information, Fig. S4

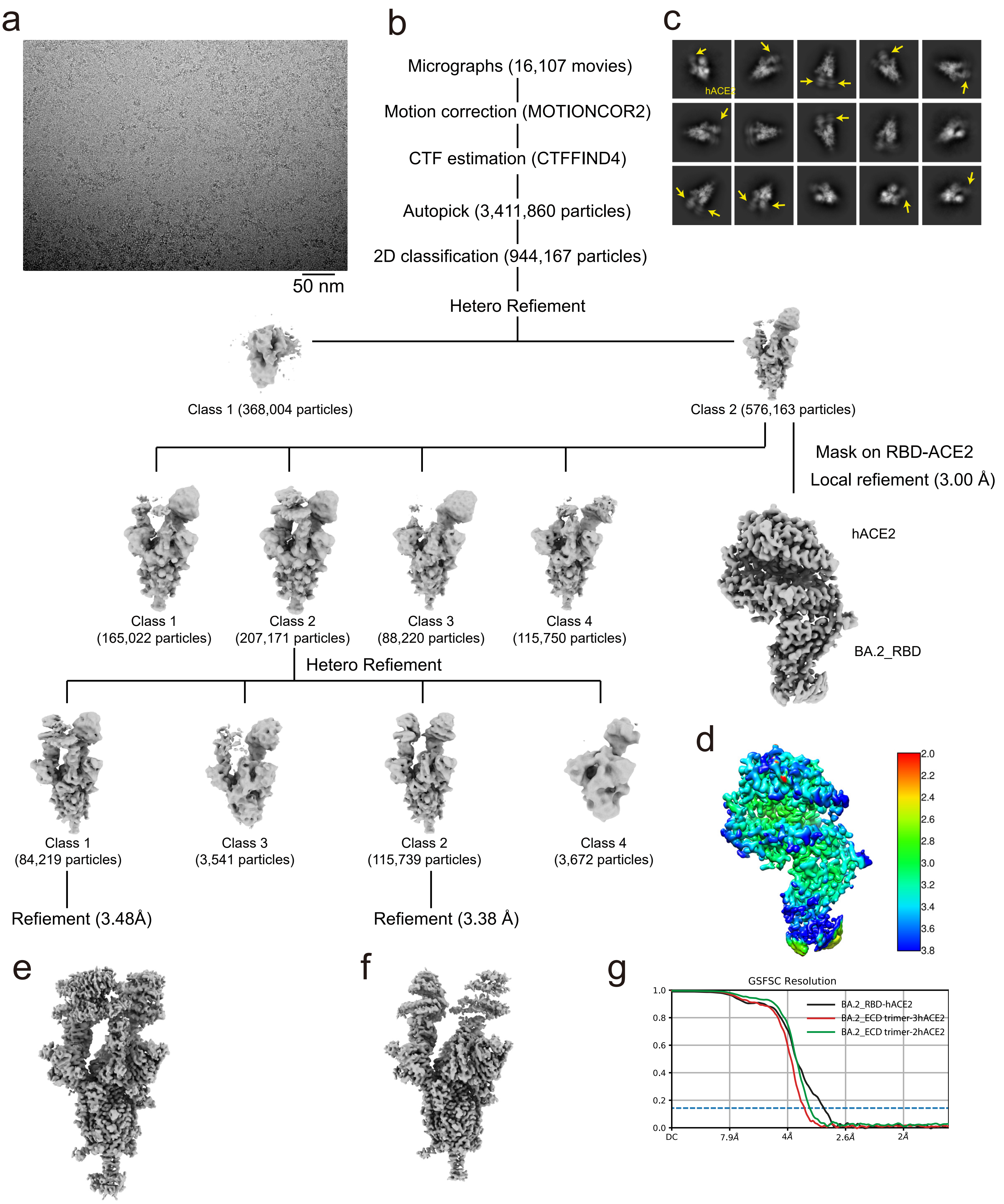

### Supplementary information, Fig. S5

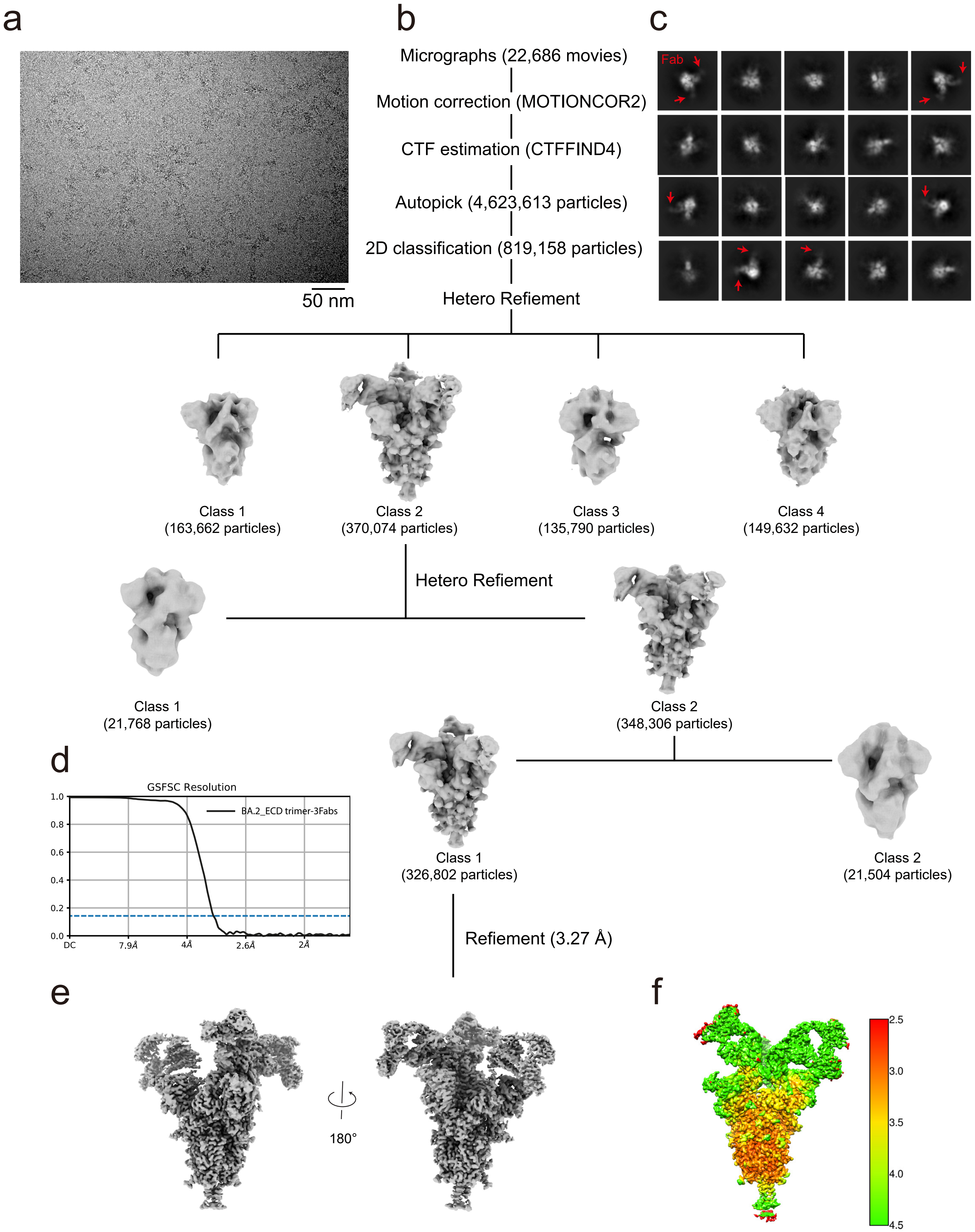

### Supplementary information, Fig. S6

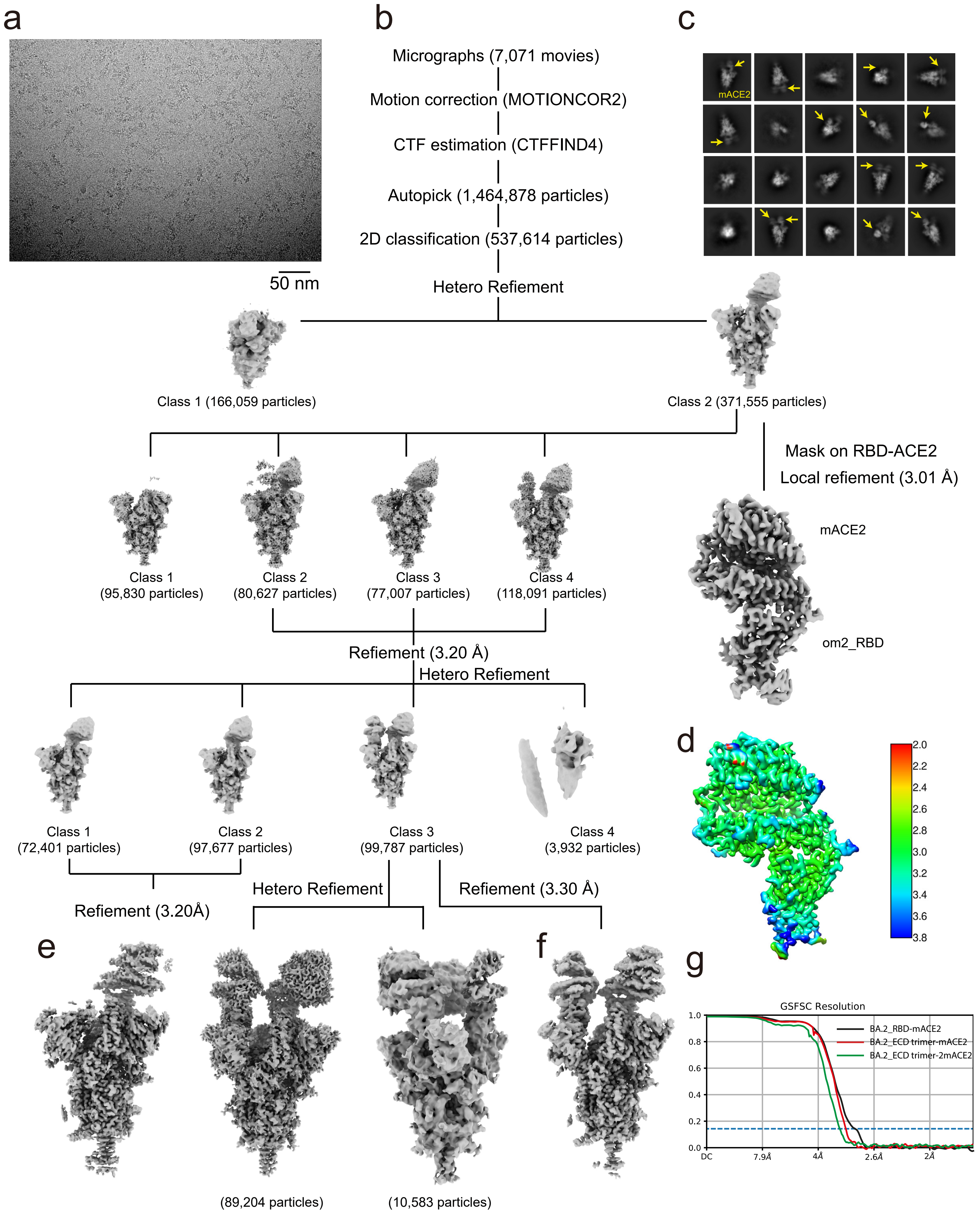

### Supplementary information, Fig. S7

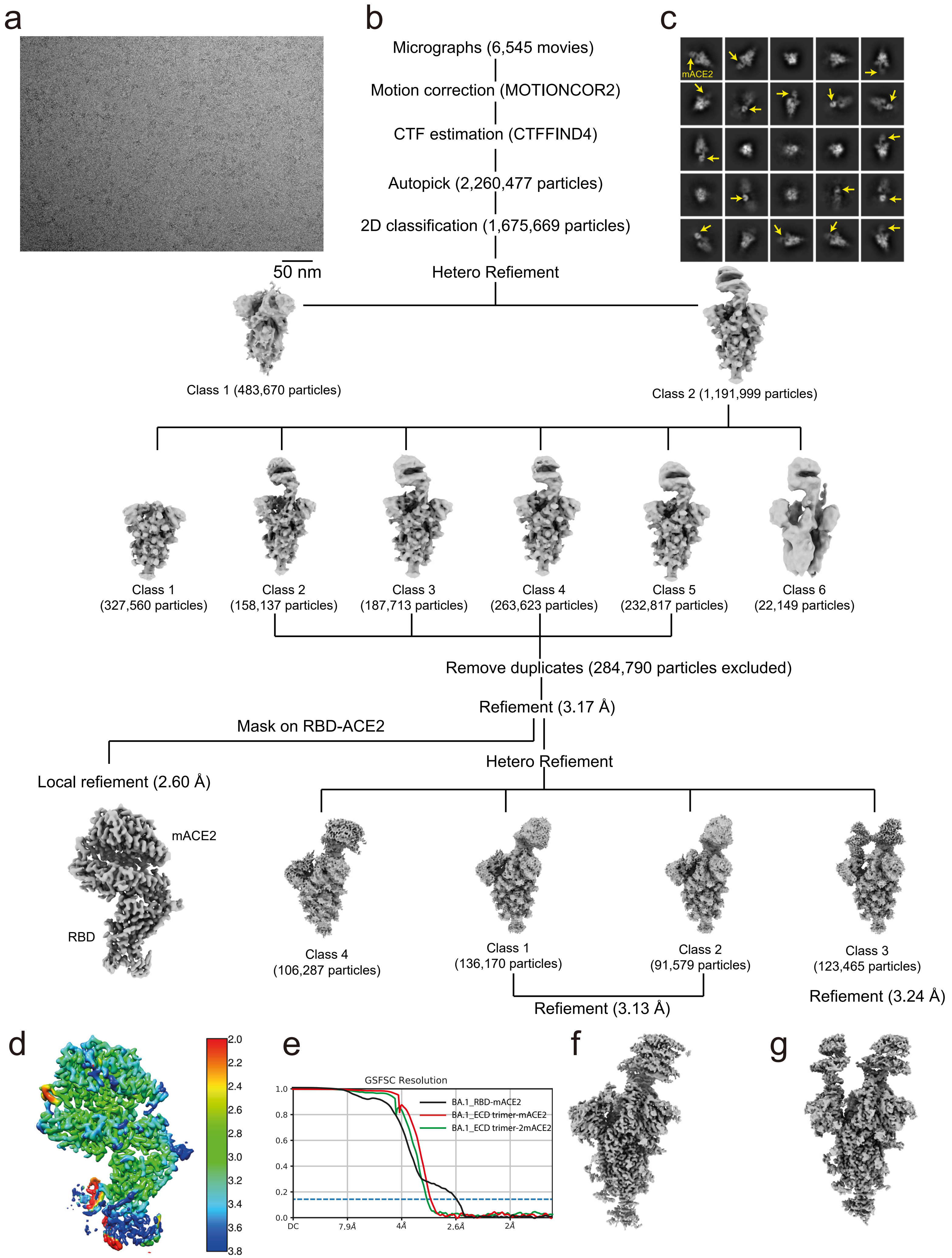

#### Materials and Methods

##### Gene cloning

The N-terminal peptidase domain (aa19-615) of ACE2 coding sequences of ACE2 from species of (human ACE2: NP _001358344.1; mouse ACE2 aa 19-615: NM _001130513.1; cat ACE2 aa 19-615: NP _001034545.1; rat ACE2: NP_001012006.1; dog ACE2 aa 19-615: NP _001158732.1; horse ACE2 aa 19-615: XP_001490241.1; sheep ACE2 aa 19-615: NP_001277036.1; pig ACE2 aa 19-615: NP_001116542.1) were optimized to *Spodoptera frugiperda* 9, and synthesized and cloned into the pFast-Bac-1 vector with an N-terminal GP67 signal peptide for secretion and a C-terminal 8×His tag. The coding sequences of the anti-Fab nanobody were cloned into the PMESy4 plasmid with an N-terminal PelB signal peptide and a C-terminal 6×His tag. All constructs were generated using Phanta Max Super-Fidelity DNA Polymerase (Vazyme Biotech Co., Ltd) and verified by DNA sequencing (Genewiz).

##### Protein expression and purification

The expression and purification of human ACE2 protein was carried out as described previously.[1] Briefly, Hi5 cells were cultured in ESF 921 serum-free medium (Expression Systems) to a density of 3 million cells/mL and then infected with baculoviruses for ACE2 at a multiplicity of infection (m.o.i.) of about 5. After 60 h of infection, the supernatant containing ACE2 was clarified by centrifugation. The secreted peptidase domain of hACE2 was captured by Ni-NTA agarose (Smart-Lifesciences) and eluted with 300 mM imidazole in HBS buffer containing 25 mM HEPES pH 7.4, 150 mM sodium chloride. The hACE2 were then purified by gel filtration chromatography using a Superdex 200 column (GE Healthcare) pre-equilibrated with HBS buffer. The fractions for hACE2 were collected, concentrated to approximately 2 mg/mL, and stored at -80 °C until use. Expression and purification of the other ACE2 proteins were performed in the same protocol as for human ACE2.

The SARS-CoV-2 spike ECD and RBD proteins were all purchased from Sino Biological Inc., including Omicron BA.1 spike ECD trimer (Cat: 40589-V08H26: containing furin cleavage site mutants and 2P mutants, i.e., R682G, R683S, R685S, K986P, and V987P), Omicron BA.2 spike ECD trimer (Cat: 40589-V08H28: with furin cleavage site mutants and 6P mutants, that is R682G, R683S, R685S, F817P, A892P, A899P, A942P, K986P and V987P), WT spike ECD trimer (Cat: 40589-V08H8: containing D614G mutants, furin cleavage site mutants and 6P mutants, that is D614G, R682G, R683S, R685S, F817P, A892P, A899P, A942P, K986P, and V987P), According to Sino Biological's description, all three ECD proteins were expressed with the bacteriophage T4 fibritin trimerization motif and a polyhistidine tag at the C-terminus.

JMB2002 is a fully human IgG1 antibody, which contains N297A mutation in its constant region to silence Fc function. JMB2002 used in this study was expressed in CHO-K1 cell line and produced by WuXi Biologics (Shanghai, China). JMB2002 Fab was transiently expressed in CHO-K1 cells and purified by Ni-NTA column via 6 x His-tag fused to the C-terminal of heavy chain (Biointron, Taizhou, China).

The anti-Fab nanobody was expressed and purified as previously described.[1] Briefly, the nanobodies were expressed in the periplasm of E. coli strain BL21(DE3) bacteria (NEB). Cultures of 2 L cells were grown to an OD600 = 0.8 at 37 °C and 180 rpm in 2×YT media containing 100 μg/mL ampicillin. Subsequently, 0.1 mM IPTG was added to the medium to induce protein expression at 28 °C and 180 rpm for an additional 8 h. Cells were harvested by centrifugation (5316 g, 30 min) and disturbed in ice-cold buffer (20 mM HEPES pH 7.4, 100 mM NaCl), then centrifuged to remove cell debris. Nb was purified by nickel affinity chromatography as previously described, followed by size exclusion chromatography using a HiLoad 16/600 Superdex 75 column. Selected fractions of Nb were finally concentrated with 10% glycerol to ~2 mg/ mL and rapidly frozen in liquid nitrogen and stored at -80 °C for further use. The quality of the purified proteins was assessed by SDS-PAGE.

**Protein complex formation**

For the ACE2 bound Omicron spike ECD protein complex, Omicron BA.1 or BA.2 spike ECD protein was incubated with the purified peptidase domain of human or mouse ACE2 at a molar ratio of 1:5 (spike trimer to peptidase domain of human ACE2) for 60 minutes on ice before purification by gel filtration chromatography using a Superose 6 increase 10/ 300 GL column (GE Healthcare) pre-equilibrated with TBS buffer (20 -mM Tris, pH8.0, 150 mM NaCl). For the JMB2002 Fab bound Omicron spike ECD protein complex, the Omicron spike ECD protein was mixed with JMB2002Fab and anti-Fab nanobody in a molar ratio of 1:5:6 (spike trimer to JMB2002Fab to anti-Fab nanobody) was incubated on ice for 60 minutes before purification by gel filtration chromatography using a Superose 6 increase 10/ 300 GL (GE Healthcare) incubated with HBS buffer (25 mM HEPES, pH 7.5, 150 mM NaCl). Fractions containing the complex were pooled and concentrated to 2 mg/ml.

**Cryo-EM data collection**

Cryo-EM grids were prepared with the Vitrobot Mark IV plunger (FEI) set to 4℃ and 100% humidity. Three-microliter of the sample was applied to the glow discharged gold R1.2/1.3 holey carbon grids. The sample was incubated for 10 s on the grids before blotting for 3 s (double-sided, blot force -2) and flash-frozen in liquid ethane immediately.

For Omicron BA.2 spike trimer-hACE2 complex, Omicron BA.2 spike trimer-JMB2002 antibody complex, Omicron BA.2 spike trimer-mACE2 complex, and Omicron BA.1 spike trimer-mACE2 complex datasets, 16107, 22686, 7071, and 6545 movies were collected respectively on a Titan Krios equipped with a Gatan K3 direct electron detection device at 300kV with a magnification of 105,000, corresponding to a pixel size 0.824 Å. Image acquisition was performed with EPU Software (FEI Eindhoven, Netherlands). We collected a total of 36 frames accumulating to a total dose of 50 $e^{-}$ $Å^{-2}$ over 2.5 s exposure.

**Cryo-EM image processing**

MotionCor2 was used to perform the frame-based motion-correction algorithm to generate drift-corrected micrograph for further processing and CTFFIND4 provided the estimation of the contrast transfer function (CTF) parameters[2, 3]. All subsequent steps including particle picking and extraction, 2D classification, three-dimensional (3D) classification, 3D refinement, and local refinement were performed using cryoSPARC[4], unless stated otherwise.

For Omicron BA.2 spike trimer-hACE2 complex dataset, a total of 3,411,860 particles were extracted from the cryo-EM micrographs. Two rounds of reference-free 2D classification, yielding 944,167 particles after clearance. Three rounds hetero refinement separated out 84,219 particles that resulted to a density of spike trimer-hACE2 structure (3:3 molar ratio) at 3.48 Å global resolution and 115,739 particles that resulted to a density of spike trimer-hACE2 structure (3:2 molar ratio) at 3.38 Å global resolution. Local refinement focused on the RBD-hACE2 with mask could reconstitute RBD-hACE2 structure at 3.00 Å global resolution.

For Omicron BA.2 spike trimer-JMB2002 antibody complex dataset, a total of 4,623,613 particles were extracted from the cryo-EM micrographs. Two rounds of reference-free 2D classification, yielding 819,158 particles after clearance. Three rounds hetero refinement separated out 326,802 particles that resulted to a density of spike trimer-Fab structure (3:3 molar ratio) at 3.27 Å global resolution.

For Omicron BA.2 spike trimer-mACE2 complex dataset, a total of 1,464,878 particles were extracted from the cryo-EM micrographs. Two rounds of reference-free 2D classification, yielding 537,614 particles after clearance. Three rounds hetero refinement separated out 170,078 particles that resulted to a density of spike bound to one mACE2 at 3.20 Å global resolution and 99,787 particles that resulted to a density of spike bound to two mACE2 at 3.30 Å global resolution. Local refinement focused on the RBD-ACE2 with mask could reconstitute BA.2 RBD-mACE2 structure at 3.01 Å global resolution.

For Omicron BA.1 spike trimer-mACE2 complex dataset, a total of 2,260,477 particles were extracted from the cryo-EM micrographs. Two rounds of reference-free 2D classification, yielding 1,675,669 particles after clearance. Three rounds hetero refinement separated out 227,749 particles that resulted to a density of spike bound to one mACE2 at 3.13 Å global resolution and 123,465 particles that resulted to a density of spike bound to two mACE2 at 3.24 Å global resolution. Local refinement focused on the RBD-mACE2 with mask could reconstitute BA.1 RBD-mACE2 structure at 2.60 Å global resolution.
Local resolution estimate was performed with cryoSPARC.

**Model building**

BA.1 spike trimer and hACE2 derived from PDB entry 7WPA structure and structure of mACE2 from Swiss-model prediction were used as the starting reference model for the building[5]. All models were fitted into the EM density map using UCSF Chimera[6] followed by iterative rounds of manual adjustment and automated rebuilding in COOT[7] and PHENIX[8], respectively. The model was finalized by rebuilding in ISOLDE[9] followed by refinement in PHENIX with torsion-angle restraints to the input model. The final model statistics were validated using Comprehensive validation (cryo-EM) in PHENIX[8]. All structural figures were prepared using Chimera[6], ChimeraX[10], and PyMOL (Schrödinger, LLC.).

**Thermal shift assay (TSA)**

The spike ECD proteins of Omicron BA.1, BA.2, and WT SARS -CoV-2 were diluted to 0.5 mg/ml with 200 mM HEPES pH 7.4, 150 mM sodium chloride (final HEPES concentration: 100 mM). Then, the diluted proteins were mixed with 10× SYPRO Orange (Thermo Fisher) and incubated for 10 min at room temperature. The reaction was performed in 384-well plates with a final volume of 10 μL. The thermal melting curves were monitored from 25°C to 80°C using a LightCycler 480 II Real-Time PCR System (Roche Diagnostics) with a ramp rate of 3.6°C per minute from 25 °C to

80 °C. Melting peaks were calculated by the LightCycler 480 software from Roche Diagnostics. A representative figure from triple experiments is shown.

**Measurement of dimeric human ACE2 binding to Omicron BA.1, Omicron BA.2, and WT SARS-CoV-2 spike ECD by biolayer interferometry**

The interaction of dimeric human ACE2 with Omicron BA.1, Omicron BA.2, and WT SARS-CoV-2 spike ECD were evaluated using Octet Red96e (Sartorius). The dimeric human ACE2 (Sino Biological, 10108-H02H) was immobilized on Protein A biosensor (Sartorius, 18-5012) followed by measuring the association and dissociation with Omicron BA.1 (Sino Biological, 40589-V08H26), Omicron BA.2 (Sino Biological, 40589-V08H28) or WT SARS-CoV-2 spike ECD protein (ACRO, SPN-C52H9). $K_D$ values were calculated with Octet Data Analysis HT 12.0 software using a 1:1 global fit model. Data were plotted using Prism V8.0 software (GraphPad). One representative figure from two independent experiments is shown.

**Measurement of anti-SARS-CoV-2 antibody JMB2002 binding to Omicron BA.2 SARS-CoV-2 spike ECD by biolayer interferometry**

The binding of anti-SARS-CoV-2 antibody JMB2002 to Omicron BA.2 spike ECD was determined using Octet Red96e (Sartorius). The biotinylation of Omicron BA.2 spike ECD protein were performed using EZ-Link Sulfo-NHS-LC-Biotin kit (ThermoFisher Scientific, A39257) following the manufacturer’s instruction. SA biosensors (Sartorius, 18-5020) were used to capture the biotinylated Omicron BA.2 spike ECD protein. The interaction of Omicron BA.2 spike ECD protein-coated sensors with different concentrations of anti-SARS-CoV-2 antibody JMB2002 were recorded prior to dissociation in kinetics buffer (0.02% Tween-20 in PBS). $K_D$ values were calculated with Octet Data Analysis HT 12.0 software using a 1:1 global fit model. Data were plotted using Prism V8.0 software (GraphPad). One representative figure from two independent experiments is shown.

**Pseudovirus neutralization assay**

In the pseudovirus neutralization assay, serial dilutions of JMB2002 IgG were preincubated with an equal volume of Omicron BA.2 pseudovirus ($2\times10^4$ $TCID_{50}$/mL) for 1 h at 37°C. Subsequently, HEK293 cells stably expressing hACE2 (Vazyme, DD1401) ($2\times10^4$ cells/well) were seeded in 96-well plates followed by the addition of pseudovirus and antibody mixtures, and incubated at 37°C for 48 h. Luciferase activity was measured using Bio-Lite Luciferase Assay System (Vazyme, DD1201). The neutralization inhibition rate was calculated using the following formula.

$$\text{Inhibition rate (\%)} = \left(1 - \frac{\text{mean intensity of sample} - \text{mean intensity of blank control}}{\text{mean intensity of negative control} - \text{mean intensity of blank control}}\right) X\,100\%$$

$IC_{50}$ values were calculated by a four-parameter logistic curve fitting approach in Prism V8.0 software (GraphPad). One representative figure from two independent experiments is shown.

**Measurement of monomeric ACE2 from different species interaction with Omicron BA.1, Omicron BA.2, and WT SARS-CoV-2 spike ECD by biolayer interferometry**

The binding of monomeric human, murine, cat, and dog ACE2 to Omicron BA.1, Omicron BA.2, and WT SARS-CoV-2 spike ECD were evaluated using Octet Red96e (Sartorius). The monomeric ACE2 of different species was biotinylated using EZ-Link Sulfo-NHS-LC-Biotin kit (ThermoFisher Scientific, A39257) following the manufacturer's instruction. SA biosensors (Sartorius, 18-5020) were then used to immobilize the biotinylated monomeric ACE2 proteins. The association and dissociation of biotinylated monomeric human, murine, cat, and dog ACE2 with different concentrations of Omicron BA.1, Omicron BA.2, or WT SARS-CoV-2 spike ECD were recorded. $K_D$ values were calculated with Octet Data Analysis HT 12.0 software using a 1:1 global fit model. Data were plotted using Prism V8.0 software (GraphPad). One representative figure from two independent experiments is shown.

#### Figure Legends for supplementary figures

**Fig. S1**. **Schematic of the Omicron BA.2 spike protein domain architecture.**

(a) Schematic of the Omicron BA.2 spike protein domain architecture. Mutations of the Omicron spike protein are labeled with different colors (blue for deleting mutation, brown for inserting mutation). Mutations in RBM are compared with WT SARS-CoV-2 and five other VOC strains. SP, signal peptide; RBM, receptor-binding motif; SD1, subdomain 1; SD2, subdomain 2; FP, fusion peptide; HR1, heptad repeat 1; HR2, heptad repeat 2; TM, transmembrane region; CT, cytoplasmic tail.

**Fig. S2**. **Binding affinities of ACE2 from different species to spike protein from different strains**

Binding affinities of ACE2 from different species to WT, BA.1 and BA.2 strains spike trimer; repetitively. There data are determined by BLI. KD values and further determined with Octet Data Analysis HT 11.0 software using a 1:1 global fit model.

**Fig. S3**. **Purification and characterization of the Omicron spike protein complex.** (a-d) Gel filtration profile of the Omicron BA.2 ECD-hACE2 complex (a), Omicron BA.2 ECD-Fab complex (b), Omicron BA.1 ECD-mACE2 complex (c), Omicron BA.2 ECD-mACE2 complex (d), all showing a sharp peak, and corresponding SDS gel showing balanced ratios for each subunit.

**Fig. S4. Cryo-EM data processing of the Omicron BA.2 spike protein-hACE2 complex.**

(a) A representative cryo-EM micrograph of Omicron BA.2 ECD-hACE2 complex with 50 nm scale bar included as a size reference. (b) Computational processing of cryo-EM data. (c) Twenty representative reference-free two-dimensional (2D) cryo-EM class averages reveal the hACE2 density (yellow arrow). (d) Local resolution of sub-reconstructions of RBD-hACE2 (left panel) and resolution bar is shown in the right panel. (e-f) The EM maps of Omicron BA.2 ECD trimer bound to 3 hACE2 (e), 2 hACE2 (f). (g)The FSC curves for the reconstructions. Color scheme: BA.2 RBDhACE2, black; BA.2 ECD trimer-3hACE2, red; BA.2 ECD-trimer-2hACE2, green. The resolution of the reconstructions using the Fourier shell cutoff at 0.143 is shown.

**Fig. S5. Cryo-EM data processing of the Omicron BA.2 ECD-Fab complex**.

(a) A representative cryo-EM micrograph of Omicron BA.2 ECD-Fab complex with 50 nm scale bar included as a size reference. (b) Computational processing of cryo-EM data. (c) Twenty representative reference-free two-dimensional (2D) cryo-EM class averages reveal the Fab density (red arrow). (d)The FSC curves for the reconstructions. (e) The EM maps of Omicron BA.2 ECD trimer-3Fabs in two front views. (f) Local resolution of sub-reconstructions of BA.2 ECD trimer-3Fabs (left panel) and resolution bar is shown in the right panel.

**Fig. S6 Cryo-EM data processing of the Omicron BA.2 spike protein-mACE2 complex.**

(a) A representative cryo-EM micrograph of Omicron BA.2 ECD-mACE2 complex with 50 nm scale bar included as a size reference. (b) Computational processing of cryo-EM data. (c) Twenty representative reference-free two-dimensional (2D) cryo-EM class averages reveal the mACE2 density (yellow arrow). (d) Local resolution of sub-reconstructions of RBD-mACE2 (left panel) and resolution bar is shown in the right panel. (e-f) The EM maps of Omicron BA.2 ECD trimer bound to 1 mACE2 (e), 2 mACE2 (f). (g)The FSC curves for the reconstructions. Color scheme: BA.2 RBD-mACE2, black; BA.2 ECD trimer-mACE2, red; BA.2 ECD-trimer-2mACE2, green. The resolution of the reconstructions using the Fourier shell cutoff at 0.143 is shown.

**Fig. S7 Cryo-EM data processing of the Omicron BA.1 spike protein-mACE2 complex.**

(a) A representative cryo-EM micrograph of Omicron BA.1 ECD-mACE2 complex with 50 nm scale bar included as a size reference. (b) Computational processing of cryo-EM data. (c) Twenty representative reference-free two-dimensional (2D) cryo-EM class averages reveal the mACE2 density (yellow arrow). (d) Local resolution of sub-reconstructions of RBD-mACE2 (left panel) and resolution bar is shown in the right panel. (e)The FSC curves for the reconstructions. Color scheme: BA.1 RBDmACE2, black; BA.1 ECD trimer-mACE2, red; BA.1 ECD-trimer-2mACE2, green. The resolution of the reconstructions using the Fourier shell cutoff at 0.143 is shown.

(f-g) The EM maps of Omicron BA.1 ECD trimer bound to 1 mACE2 (f), 2 mACE2 (g).
